# Multi-hit autism genomic architecture evidenced from consanguineous families with involvement of FEZF2 and mutations in high-risk genes

**DOI:** 10.1101/759480

**Authors:** Mounia Bensaid, Yann Loe-Mie, Aude-Marie Lepagnol-Bestel, Wenqi Han, Gabriel Santpere, Thomas Klarić, Christine Plançon, Rim Roudies, Imane Benhima, Nisrine Tarik, Patrick Nitschke, Jean-Marc Plaza, Hassan Kisra, Jean-François Deleuze, Nenad Sestan, Jean-Marie Moalic, Michel Simonneau

**Affiliations:** Hôpital Militaire d’Instruction Mohammed V. Rabat, Maroc; Centre Psychiatrie et Neurosciences, INSERM. Paris, France; Departments of Neuroscience of Genetics, of Psychiatry, and of Comparative Medicine, Program in Cellular Neuroscience, Neurodegeneration and Repair, and Yale Child Study Center, Yale School of Medicine, New Haven, CT 06510, USA; Centre National de Genotypage, CEA. Evry, France; Centre de Pédopsychiatrie CHU Arrazi, Salé, Maroc; Faculté de Médecine Site Necker. Paris, France; LAC-CNRS, 91400 Orsay, France; Département de Biologie, Ecole Normale Supérieure Paris-Saclay, Université Paris-Saclay, France

**Author notes:** The first three authors should be regarded as joint First Authors.

## Abstract

Autism Spectrum Disorders (ASDs) are a heterogeneous collection of neurodevelopmental disorders with a strong genetic basis. Recent studies identified that a single hit of either a *de novo* or transmitted gene-disrupting, or likely gene-disrupting, mutation in a subset of 65 strongly associated genes can be sufficient to generate an ASD phenotype. We took advantage of consanguineous families with an ASD proband to evaluate this model. By a genome-wide homozygosity mapping of ten families with eleven children displaying ASD, we identified a linkage region of 133 kb in five families at the 3p14.2 locus that includes *FEZF2* with a LOD score of 5.8 suggesting a founder effect. Sequencing *FEZF2* revealed a common deletion of four codons. However, the damaging *FEZF2* mutation did not appear to be sufficient to induce the disease as non-affected parents also carry the mutation and, similarly, *Fezf2* knockout mouse embryos electroporated with the mutant human *FEZF2* construct did not display any obvious defects in the corticospinal tract, a pathway whose development depends on *FEZF2*. We extended the genetic analysis of these five *FEZF2*-linked families versus five *FEZF2* non-linked families by studying *de novo* and transmitted copy number variation (CNV) and performing Whole Exome Sequencing (WES). We identified damaging mutations in the subset of 65 genes strongly associated with ASD whose co-expression analysis suggests an impact on the prefrontal cortex during the mid-fetal periods. From these results, we propose that both *FEZF2* deletion and multiple hits in the repertoire of these 65 genes are necessary to generate an ASD phenotype.

**Significance Statement:** The human neocortex is a highly organized laminar structure with neuron positioning and identity of deep-layer cortical neurons that depend on key transcription factors, such as *FEZF2, SATB2, TSHZ3* and *TBR1*. These genes have a specific spatio-temporal pattern of expression in human midfetal deep cortical projection neurons and display mutations in patients with Autism Spectrum Disorder (ASD). Here, we identified a linkage region involving FEZF2 gene in five consanguineous families with an ASD proband. For these FEZF2-allele linked probands, we identified a four-codon deletion in FEZF2 and damaging mutations in other high-risk ASD genes, that exhibit regional and cell type–specific convergence in neocortical deep-layer excitatory neurons, suggesting a multi-hit genomic architecture of ASD in these consanguineous families.

## Introduction

The genetics and genomics of neuropsychiatric disorders (NPDs) have made unprecedented progress recently, in particular for unravelling the genetic architecture of Autism Spectrum Disorders (ASDs) (1, 2). ASD is a phenotypically and genetically heterogeneous neurodevelopmental condition that is characterized by deficits in reciprocal social interaction, repetitive behavior patterns, and restricted interests (3), with a prevalence as high as 1 in 68 children (4). Both *de novo* and transmitted multiple definitive risk-carrying copy number variations (CNVs), protein-altering mutations, and/or non-coding alleles have been identified in ASD (5, 6). Recent studies leveraged the statistical power afforded by rare *de novo* putatively damaging variants and have identified more than 65 strongly associated or high-confidence genes (5). The most deleterious variants that are likely gene-disrupting in the highest confidence subset (False Discovery Rate, FDR ≤ 0.01) of these genes (n = 30) as a group confer ∼20-fold increase in risk (5). To date, there is very little evidence for a multi-hit hypothesis vis-á-vis these highly damaging *de novo* coding mutations (6). Willsey et al. proposed a model with a single highly protein-disrupting mutation causing functional haploinsufficiency and thus sufficient to confer these very large risks (6).

Our aim was to study consanguineous families with autism probands in order to test whether the single hit model applies to these families with a possible founder effect, and whether single hit high-risk genes are part of these 65 strongly associated genes.

Homozygosity mapping in pedigrees with shared ancestry has been a powerful methodology to identify autosomal recessive disease genes in ASDs and to advance our understanding of the genetic architecture of these complex diseases (7, 8, 9). Here, we identified ten simplex, consanguineous families from Morocco and performed whole-genome linkage studies and whole-exome sequencing (WES) in order to validate a genetic architecture based either on a single hit model or on multiple hits.

## Results

### Homozygosity mapping implicates FEZF2-linked locus in ASD etiology

We used homozygosity mapping, CNV analysis followed by WES, and Sanger sequencing strategies to identify disease-related mutations in ten trios (parents and affected child) from consanguineous ASD families.

We first performed a genome-wide homozygosity mapping linkage analysis in ten trios in which the parents are either first or second cousins. All patients were examined and diagnosed using ADI-R (**Supplementary Table S1**). The M/F ratio of affected children in these consanguineous families was 2.6:1 (8 males, 3 females), and this ratio is comparable to the ratio found in consanguineous multiplex pedigrees from a previous study (8). The linkage analysis revealed several genetic loci overlapping between pedigrees with strong support for linkage with logarithm of odds (LOD) scores ranging from 3.6 to 5.8 (**Supplementary Table S2**). One region of 133 kb that includes the *FEZF2* gene at 3p14.2 was identified in five families (2, 5, 6, 18, 23) and had a higher LOD score of 5.8 (**Figure 1A-B**). The *FEZF2* gene encodes the transcription factor required for the specification of corticospinal motor neurons (10, 11, 12). Altogether, these results suggest that a founder locus that includes *FEZF2* is associated with ASD in five families.

**Figure 1.**
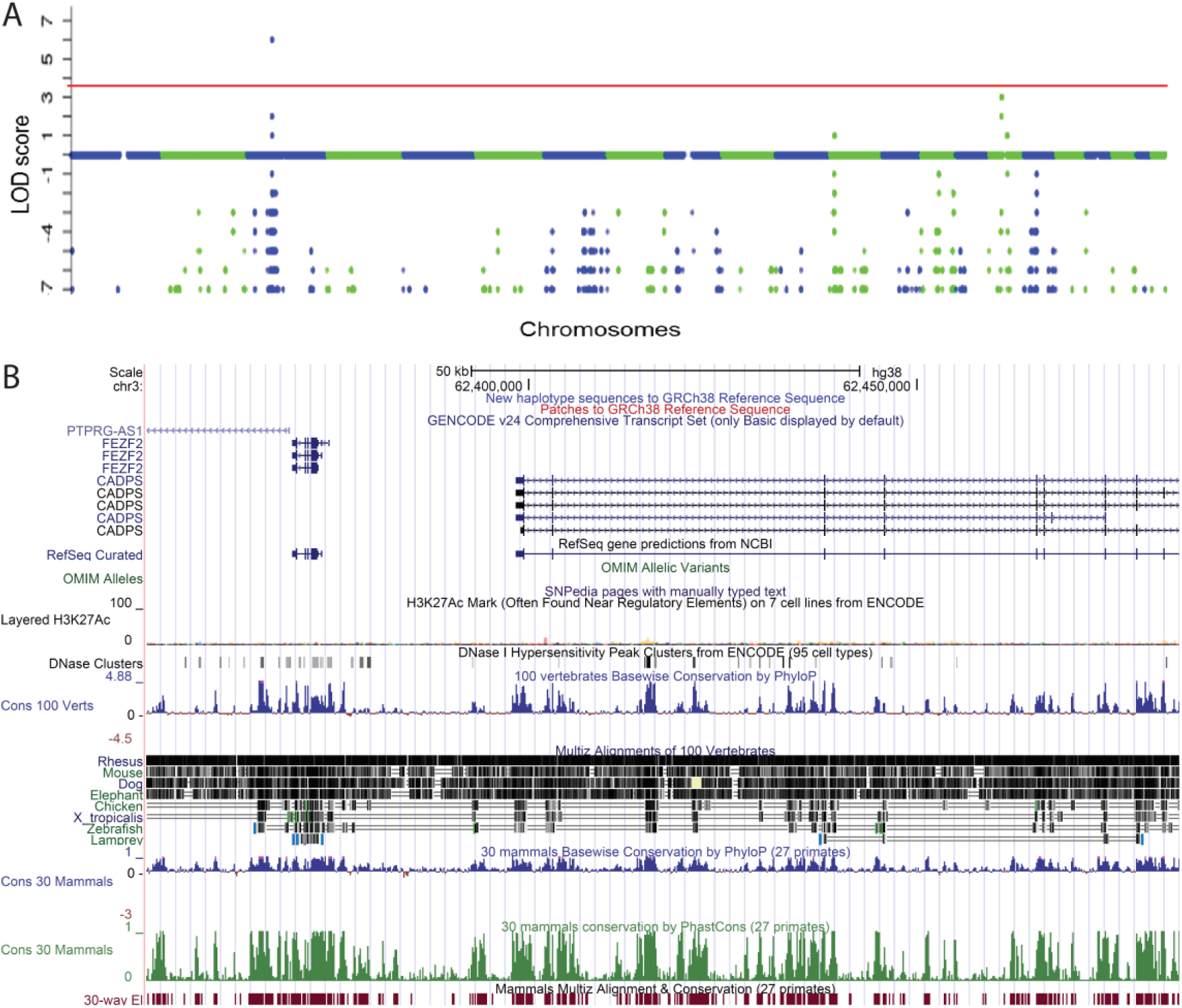
The FEZF2-linked locus as a founder locus. (**A**) Homozygosity mapping linkage results. The Manhattan plots display the LOD scores of each analyzed SNP marker along families 2, 5, 6, 18, 23. Alternating blue and green colors indicate changes in chromosomes (1 to 22), red line indicates the significance threshold (LOD score at 3.6). Significant Linkage observed in chromosome 3 at region 3p14.1 and in chromosome 16 at 16p11.2. (**B**) The 133kb region at 3p14.2 visualized in the context of the genome using the UCSC Browser.

*FEZF2* sequencing revealed heterozygous deletions of 12bp in exon 2 (**Figure 2A**). These mutations, either transmitted by one parent (father) for patients 32, 318 and 418 or arising *de novo* for patients 36 and 323, lead to an in-frame deletion of 4 amino acids (RLPA) (Arg237_Ala240del) in the FEZF2 protein (**Figure 2B**). This core region showed mammalian conservation (**Figure 2B**). Interestingly, haplotype analysis indicated that this *FEZF2* deletion was found in different haplotypes (**Figure 2C**). In our study, both transmitted and *de novo FEZF2* mutations (n=4) were found in haplotype SQ4 that is linked to ASD in this group of families. In haplotype SQ1, which is the most frequent, we identified one individual from Colombia with the same RPLA mutation out of 2,535 individuals from the 1,000 genome database (http://www.internationalgenome.org/). A similar mutation was found in three patients with intellectual disability out of 2,379 screened Saudis and was also found as homozygous in one autistic behavior patient (13). Their haplotype is either haplotype SQ4, SQ18 or SQ20 (**Supplementary Table S3**). The SQ1 (68.7%), SQ2 (8.4%) and SQ4 (5.8%) haplotypes differ by one, two and three mutational events, respectively. Multidimensional scaling (MDS) analysis of these FEZF2 locus haplotypes from the 1,000 genomes data indicated that the FEZF2 alleles from Moroccan families are related to the African FEZF2 allele (**Figure 2D**). As for other mutations found in brain diseases (14), one can propose that the mutation arose independently in each case.

**Figure 2.**
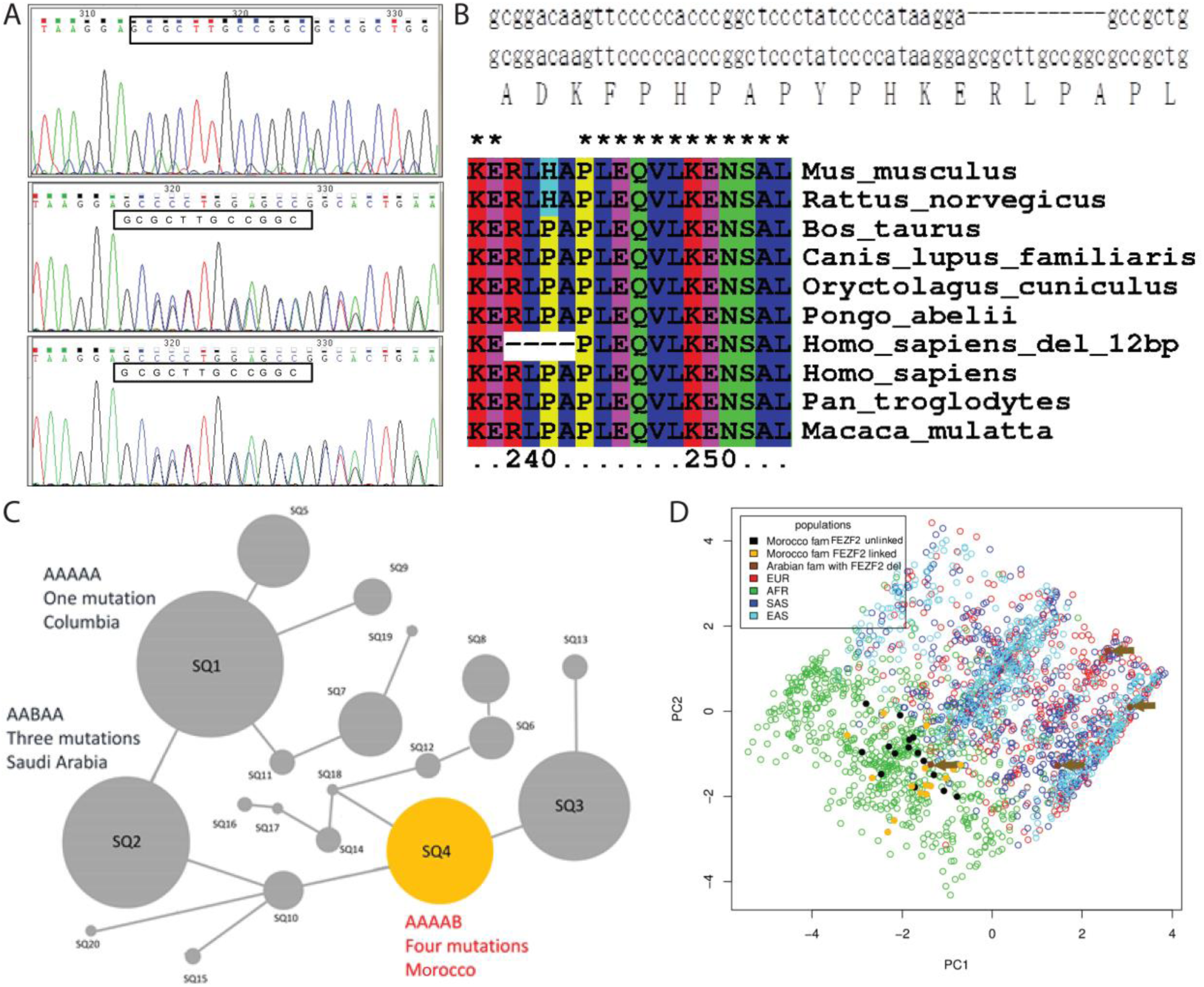
Identification of a mutation in the coding sequence of the FEZF2 gene. (**A**) Heterozygous deletion of 12bp in FEZF2 exon 2. Top panel: mother, middle panel: father, lower panel: autistic child. (**B**) The in-frame mutation results in deletion of 4 amino acids (RLPA) (Arg237_Ala240del). Sequence alignments of the affected FEZF2 region showed mammalian conservation. (**C**) Haplotype network of the FEZF2 RLPA core region in our multi-ethnic panel from the 1,000 genomes data showing Saudi Arabian patients and our Moroccan FEZF2 allele-linked ASD families. Each circle represents a haplotype, the diameter of which is proportional to the haplotype frequency in the sample. Each line connecting two haplotypes represents a mutational event. The RLPA mutation was found in the SQ4 (Morocco patients), SQ2 (Saudi Arabia patients) and SQ1 (Columbian person) haplotypes. These three haplotypes differ by one to three mutational events, suggesting a possible unique founding origin of the RLPA mutation in FEZF2. (**D**) Two-dimensional plot of multidimensional scaling (MDS) analysis of FEZF2 locus haplotypes from the 1,000 genomes data. The FEZF2 alleles from Moroccan families are related to the African FEZF2 allele. Similarly, one of the parents from Saudi Arabia displayed a FEZF2 allele similar to African FEZF2 alleles.

Brain MRI with and without contrast injection was performed for the autistic twins, 318 and 418 of Family 18, and for patient 32 of Family 2 (see **Supplementary Table S1**). The MRI results did not show any significant abnormalities, and no cerebral atrophy or significant cortical dysplasia were detected. Interestingly, the corpus callosum did not display any significant abnormalities.

### Functional analysis of the mutant FEZF2 protein *in vivo* in the context of neocortical neuronal projection during embryogenesis

The *FEZF2* gene (also known as FEZL, ZF312 or ZNF312) regulates the molecular identity, dendritic development and axon targeting of layer 5 corticospinal neurons in the cerebral cortex (10, 11, 12). TBR1 directly binds to FEZF2 regulatory regions and represses its activity in Layer 6 cortico-thalamic projection neurons to restrict the origin of the corticospinal tract to layer 5 (15, 16, 17, 18). TBR1 is one of the 20 ASD high risk candidates that confer ∼20-fold increase in risk (5). Furthermore, cis-regulatory regions controlling the expression of *FEZF2* and its targets, such as PLEX1, display evolutionarily conserved roles in the formation of direct corticospinal connections in primates (19, 20).

Since the FEZF2 protein is conserved from mouse to human (Percent identity: 95.22% and Percent similarity: 95.43%) and the *FEZF2* mutation involved conserved amino acids (RLPA amino acids), we tested the function of the mutated protein in a mouse conditional *Fezf2* knockout background (**Supplementary Table S4**).. Electroporation of either the wild-type or mutant human *FEZF2* expression constructs resulted in normal corticospinal tract projections emanating from layer 5 (**Figure 3A**) indicating that expression of the FEZF2 protein with the RPLA deletion is not sufficient to impact the long-distance axonal projection that characterizes FEZF2-expressing cortical neurons from Layers 5 (**Figure 3B**). Given that FEZF2 regulates the fate choice of subcortical projection neurons in the developing cerebral cortex (15), we next assessed the function of the mutant FEZF2 protein in corticospinal tract development. Using the same model as outlined above, we evidenced GFP expression in pons of electroporated embryos that expressed the mutant FEZF2 protein similar to that previously reported in control animals (10). These results suggest that the mutant FEZF2 protein retains its normal function, at least in the context of corticospinal tract development and dendrite maturation, which is consistent with the observation that the RPLA FEZF2 mutation can be found in unaffected parents (family 2 and family 18) and is therefore not sufficient to induce an ASD phenotype (**Figure 3C**).

**Figure 3.**
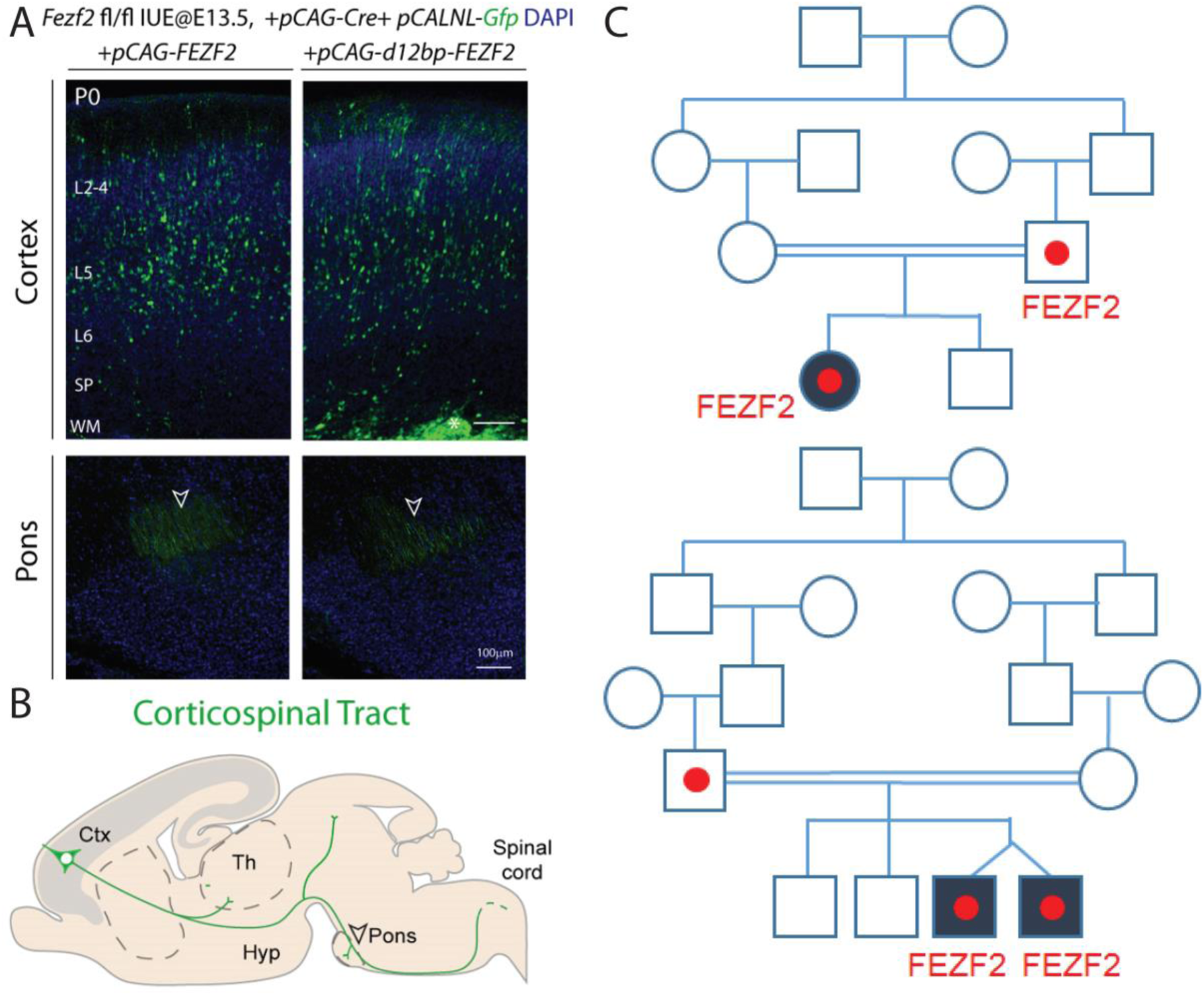
The mutant version of human *FEZF2* rescued the corticospinal tract defects associated with the loss of mouse *Fezf2* in layer 5 neurons. (**A**) Schematic representation of corticospinal projections originating from deep layer cortical neurons (Layers 5). Abbreviations: Ctx: Cortex; Th: Thalamus; Hyp: Hypothalamus. (**B**) E13.5 Fezf2^fl/fl^ mouse embryos were electroporated *in utero* with a combination of three expression constructs; pCAG-Cre to excise the endogenous mouse Fezf2 gene, pCALNL-GFP to visualize transfected cells, and either pCAG-FEZF2 to express the wildtype human FEZF2 protein or pCAG-d12bp-FEZF2 to express the mutant FEZF2 protein carrying the 12bp deletion. Electroporations were targeted to layer 5 of the cortex (upper panels) and at P0, in both groups, labelled axons were seen projecting to the pons indicating normal corticospinal tract development. The *in utero* electroporations were done in the same liter, on different embryos, targetting the left hemisphere with wildtype human FEZF2 and the right hemisphere (of a different embryo) with the 12bp-FEZF2. Mean axonal length of 108.17±40.38 at P0, for 3 animals with RPLA deleted *FEZF2* transgene expression; n=25 neurons analyzed. (**C**) Consanguineous pedigrees with ASD in two FEZF2-linked families. Deletion of 12bp in the FEZF2 gene occurs in two parents with transmission to probands. Squares indicate males and circles indicate females. Individuals affected with ASD are presented as black symbols, white symbols are unaffected. Symbols containing a • indicate carriers of the 12bp deletion in the FEZF2 gene.

### Identification of *de novo* Copy Number Variants (CNVs) and Single Nucleotide Variants (SNVs)

We first analyzed *de novo* CNVs in siblings from both *FEZF2*-allele linked and unlinked families (**Supplementary Table S5**). We found 56 and 21 *de novo* CNVs in *FEZF2*-allele unlinked and linked families, respectively. This number of *de novo* CNVs in our cohort (found for 11 of 11 patients) is considerably higher than the previously reported number for non-consanguineous families (21). We analyzed these *de novo* CNVs using Disease Association Protein-Protein Link Evaluator (DAPPLE) to study the gene networks involved. DAPPLE looks for significant physical connectivity among proteins encoded by genes in loci associated with disease according to protein-protein interactions reported in the literature (22, 23). For *de novo* CNVs from *FEZF2*-unlinked families (N= 56), the DAPPLE network is statistically significant for indirect connections (Seed Indirect Degrees Mean, P = 7.99 × 10 ^−3^ and Common Interactors Degrees Mean P = 3.99 × 10^−2^) (**Figure 4A**). In contrast, for *de novo* CNVs from *FEZF2*-linked families (N=21), no significant network was found using DAPPLE. Interestingly, for the DAPPLE network of *FEZF2* allele-unlinked families, nodes of the gene network involve genes encoding presynaptic proteins, synaptic receptor subunits, and adhesion molecules: PTPRD, NSF (24, 25), GRM7, GRIK2 (26), GABRG1 (27), ROBO2 (28), EPHA7, and ALCAM (29). *DUSP22*, a gene that encodes a protein activating the Jnk signaling pathway, was already found to be mutated in another ASD study (30). To complement the DAPPLE analysis, we used Over-Representation Analysis (ORA) from WebGestalt suite (31). ORA identified a network that includes GRM7 and GRIK2, genes that encode synaptic proteins, thus implicating the involvement of a GO:0007215 glutamate receptor signaling pathway (Ontology: biological_process) with p=2.22e-16 (Cellular Component: GO:0045211: postsynaptic membrane; p <2.22e-16; Molecular Function: GO:0008066: glutamate receptor activity; p <2.22e-16) (**Figure 4B**).

**Figure 4.**
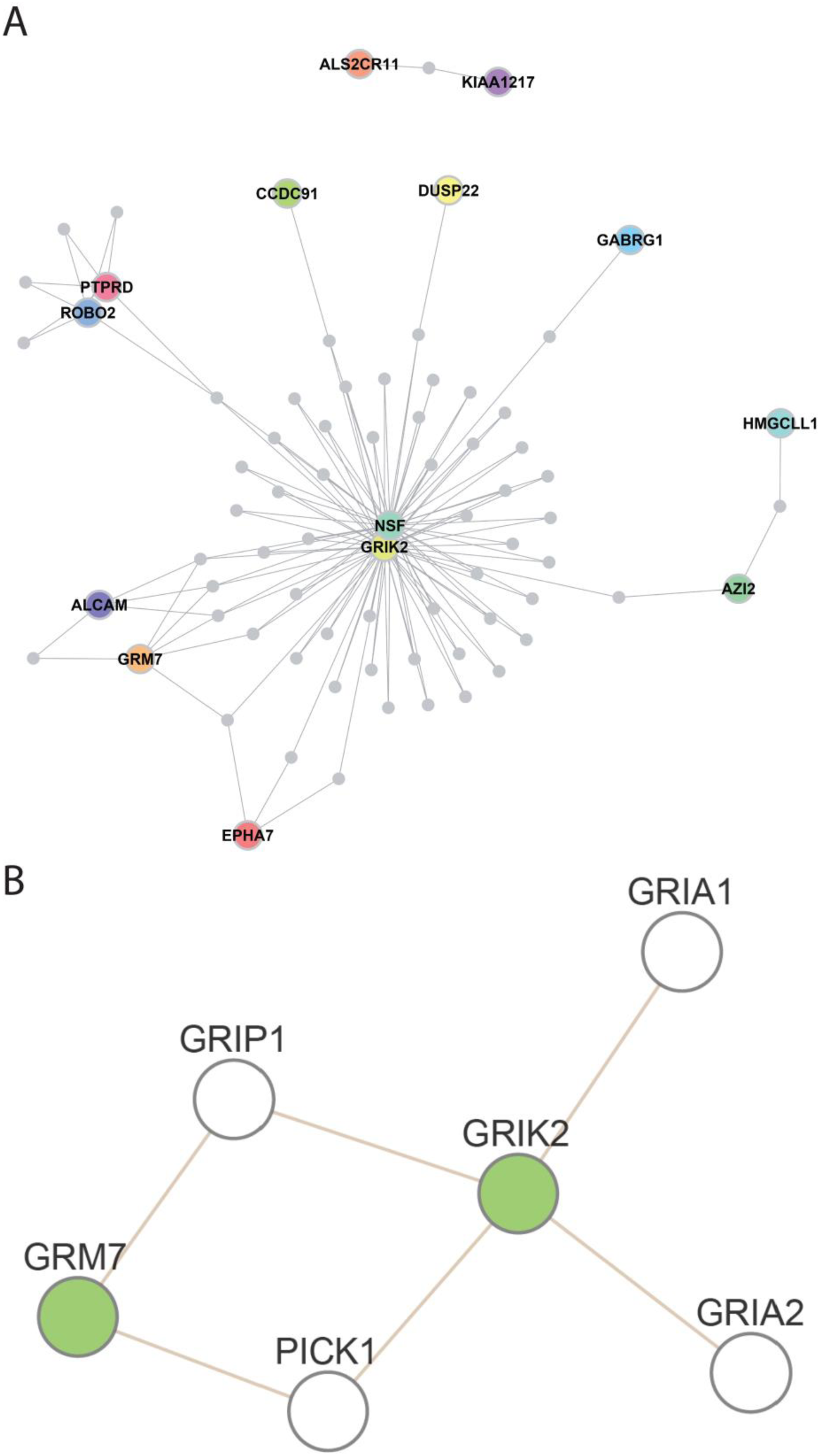
Interactome network from products of *de novo* CNV genes found in FEZF2-unlinked families. (**A**) Disease Association Protein-Protein Link Evaluator (DAPPLE) network derived from gene hits found in *de novo* CNVs. The network is statistically significant for indirect connections. (**B**) Webgestalt analysis of the same *de novo* CNVs demonstrating a GO:0007215 glutamate receptor signaling pathway (Ontology biological_process) with p=2.22e-16.

We next performed whole exome sequencing (WES) in ten trios and confirmed by Sanger sequencing eleven *de novo* mutations (**Supplementary Table S6**). We found *de novo* single nucleotide variants (SNVs) in *ERN1, HADHB, DMKN*, and *STRIP2* in three *FEZF2-*allele linked patients and variants in *SH3TC2, CDCA4, APOA, PHCTR3, Elf4G1, GRK4*, and *TMEM11* were found in five patients from *FEZF2* allele-unlinked families. All variants were novel and not observed in the exome variant server (EVS, http://evs.gs.washington.edu/EVS/). These variants impact amino acids conserved in mammals. This ratio of 11 protein-coding *de novo* variants in our study is rather similar to that described for intellectual disability cases with *de novo* variants resulting in a total of 84 *de novo* mutations in the protein-coding regions for 50 patients (32).

### Analysis of *de novo* and transmitted CNVs and SNVs in FEZF2 allele-linked and unlinked families

Each person inherits mutations from their parents (transmitted mutations), some of which may predispose the person to certain diseases. Meanwhile, new mutations may occur spontaneously during the reproductive process (*de novo* mutations) and, if disrupting key genes, may similarly increase the risks of disease. It is therefore important to study the combination of *de novo* and transmitted mutations.

We first performed this analysis for *de novo* and transmitted CNVs in these 10 consanguineous families and we found 337 and 314 genes for CNVs of *FEZF2* allele-unlinked and linked families, respectively (**Supplementary Table S5**). We used GO to analyze the gene networks involved. In *FEZF2* allele-linked families, we identified enrichment in GO: 0007156 biological process: homophilic cell adhesion via plasma membrane adhesion molecules and GO: 0050911: detection of chemical stimulus involved in sensory perception of smell classes with P=2.54 E^−06^ and P=1.04 E^−06^, respectively. In *FEZF2* allele-unlinked families, we similarly found the involvement of GO biological process GO:0007156: homophilic cell adhesion via plasma membrane adhesion molecules with P=9.25E^−08^. Furthermore, DAPPLE analysis identified statistically significant networks in FEZF2 allele-linked families (**Supplementary Figure 1A**) and FEZF2 allele-unlinked families (**Supplementary Figure 1B**). For the *FEZF2* allele-linked families, the DAPPLE network is significant for both direct and indirect nodes (Direct Edges Count P = 1.99 × 10^−2^; Seed Direct Degrees Mean P = 3.99 × 10^−3^; CI Degrees Mean P = 5.99 × 10^−3^). For the *FEZF2* allele-unlinked families, the DAPPLE network is also significant for both direct and indirect nodes (Direct Edges Count P = 3.49 × 10^−2^; Seed Indirect Degrees Mean P = 1.99 × 10^−2^; CI Degrees Mean P=1.99 × 10^−3^). Interestingly, mutations found in *de novo* CNVs of *FEZF2* allele-unlinked families underline the importance of synaptic networks as already reported (5, 6, 33–35). In contrast, results from transmitted and *de novo* CNVs of *FEZF2* allele-linked and unlinked families point to distinct gene classes with adhesion proteins involved in hemophilic interactions, such as clustered protocadherins (36, 37), that are known to be involved in the establishment of the diversity of neural circuit assemblies and olfaction pathways. Links between olfaction and autism were recently found in ASD patients (38, 39). From these CNV data (**Supplementary Table S5**), we also analyzed genes repeatedly found mutated in different families. We identified 207 genes mutated in more than one family (from 2 to 10 families) (**Supplementary Table S7**). Interestingly, their analysis by Gene Ontology found enrichment in genes of GO: 0007156∼homophilic cell adhesion via plasma membrane adhesion molecules (P=1.19E^−7^), GO: 0098742∼cell-cell adhesion via plasma-membrane adhesion molecules (P=2.52E^−7^) and GO: 0050911∼ detection of chemical stimulus involved in sensory perception of smell (P=1.53E^−9^).

We also investigated transmitted SNVs mutations and identified 974 and 931 mutated genes in *FEZF2* allele-linked and unlinked families, respectively, with the intersection of the two groups containing 211 genes (**Supplementary Table S8**). Transmitted SNVs for *FEZF2* allele-linked families (N=974) display enrichment in genes of GO biological process GO:0021795 cerebral cortex cell migration (P=2.04E^−05^) and GO:0021885 forebrain cell migration (P=1.54E^−05^). In contrast, transmitted SNVs for *FEZF2* allele-unlinked families (N=931) display enrichment in genes of GO biological process GO:0007155 cell adhesion (P=1.52E^−06^).

### High-risk ASD genes involvement for *de novo* and transmitted CNVs and SNVs in both FEZF2 allele-linked and unlinked families

Studies leveraging the statistical power afforded by rare *de novo* putatively damaging variants have identified more than 65 genes strongly associated with ASD (5, 35). The most deleterious variants in the highest confidence subset of these genes (n = 30) as a group confer ∼20-fold increase in risk (5). In four *FEZF2* allele-linked families, we identified transmitted SNV mutations in eight genes strongly associated with ASD: *KAT2B*, *DIP2A, NINL, MFRP, AKAP9, ASH1L, SCN2A*, and *ZNF559* (**Figure 5**). In two *FEZF2* allele-linked families (Family 6 and 23), the *FEZF2* mutation is a *de novo* mutation. In five *FEZF2* allele-unlinked families, we identified transmitted SNVs and CNV in six genes strongly associated with ASD: *APH1A*, PTK7, *CTTNBP2, CHD8, SETD5*, and *KATNAL2* (**Figure 6**). For both *FEZF2* allele-linked and -unlinked families, we identified an enrichment of mutations in genes strongly associated with ASD (2.16 fold compared to expectations with a hypergeometric P value = 4.22 × 10 ^−3^). Interestingly, from the 14 mutations identified in high-risk ASD genes, thirteen are found in transmitted SNVs and only one in a transmitted CNV (CHD8).

**Figure 5.**
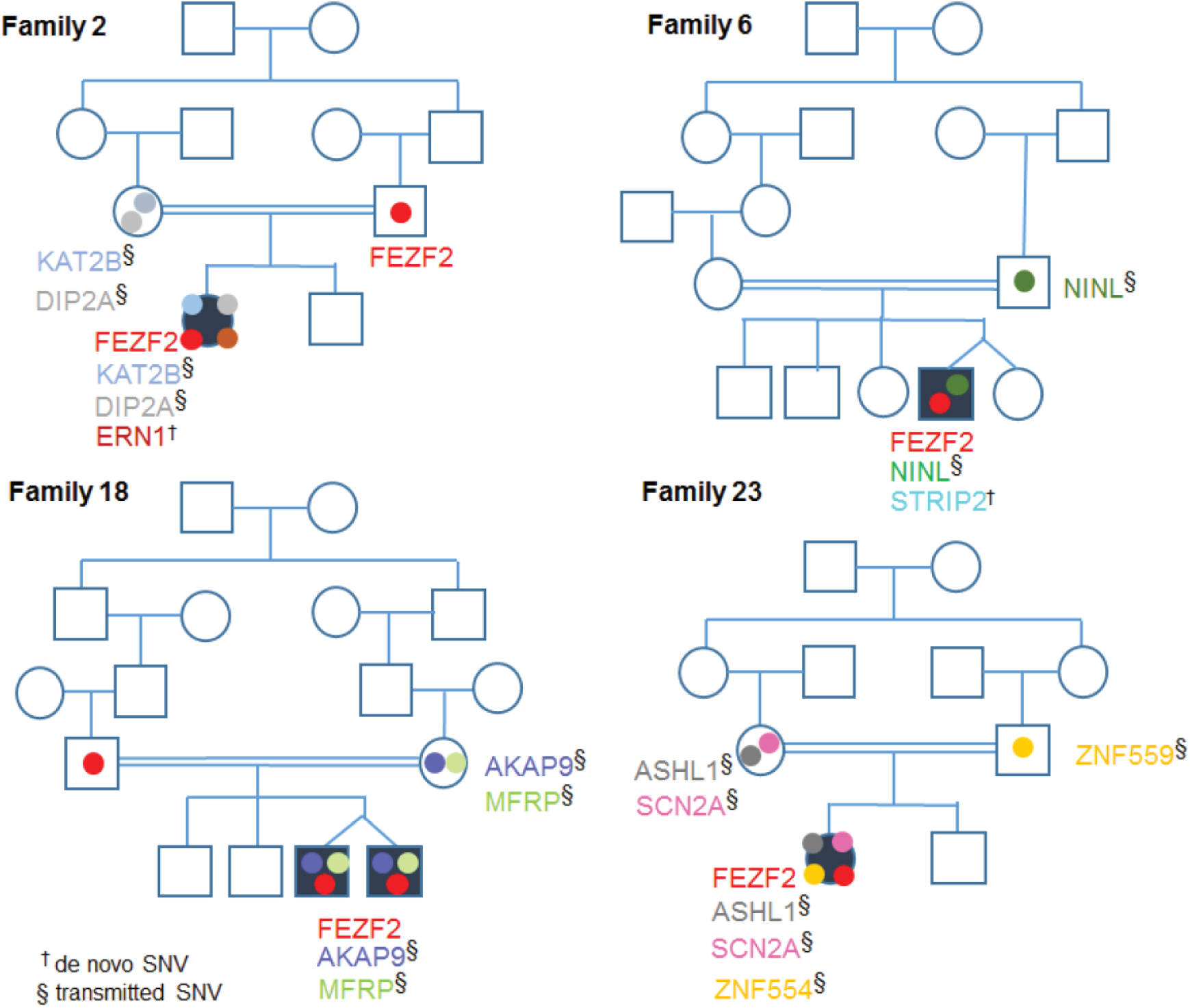
Multiple hits in ASD risk genes are necessary to generate an ASD phenotype. Consanguineous pedigrees with ASD in four FEZF2 allele-linked families: Family 2 whose autistic child displayed the transmitted mutations in the *FEZF2, DIP2A*, and *KAT2B* genes as well as the *de novo* SNV in the *ERN1* gene. Family 6 whose autistic child displayed *de novo* mutations in the *FEZF2* and *STRIP2* genes as well as a transmitted mutation in *NINL* gene. Family 18 whose twin autistic children displayed transmitted mutations in the *FEZF2, AKAP9* and *MFRP* genes. Family 23 whose autistic child displayed the *de novo* mutation in *FEZF2* and transmitted mutations in the *ASH1L, SCN2A* and *ZNF559* genes. Squares indicate males and circles indicate females. Individuals affected with ASD are presented as black symbols, white symbols are unaffected. †de NOVO SNV, § transmitted SNV

**Figure 6.**
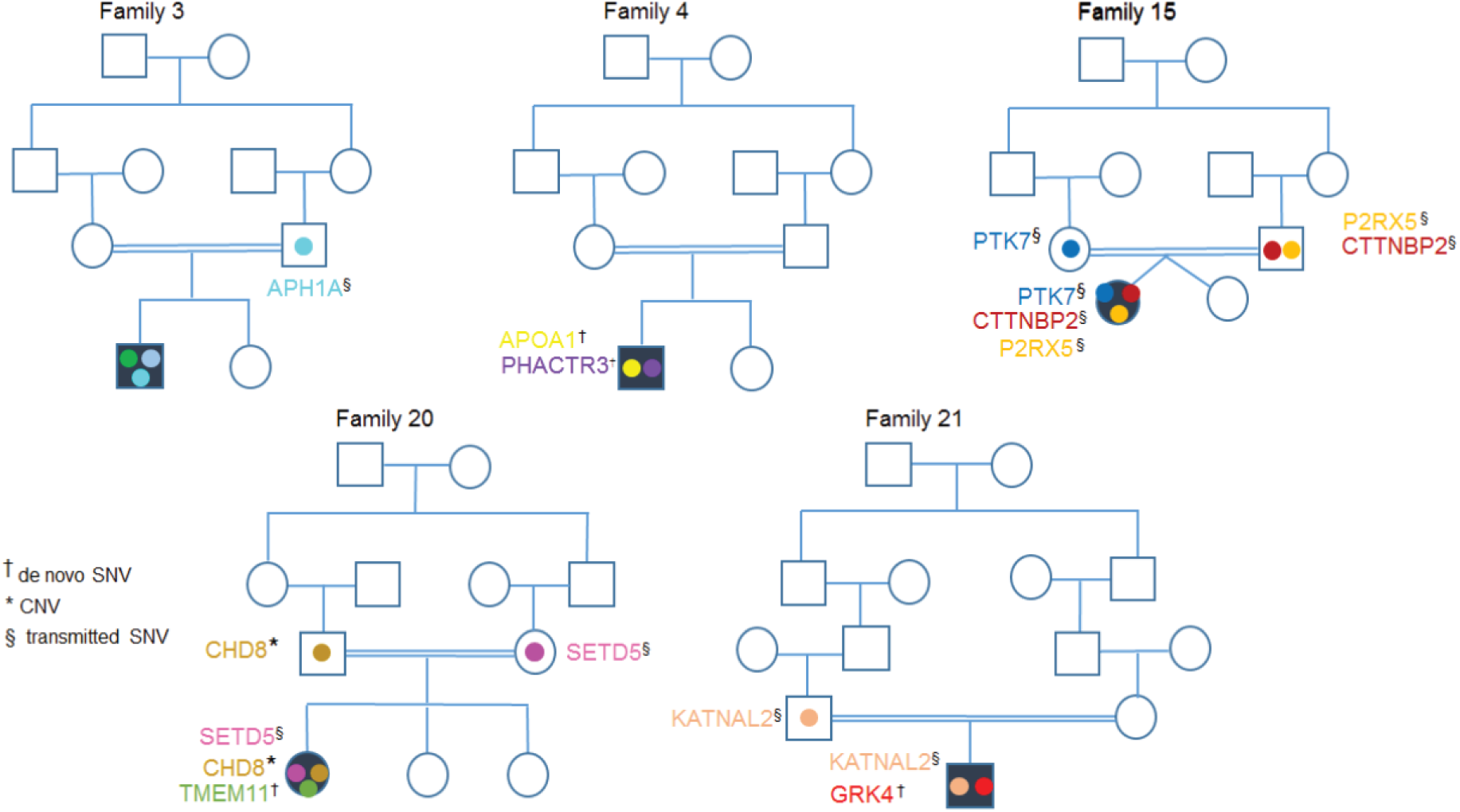
Multiple damaging mutations in the high-risk genes in FEZF2 allele-unlinked families. Family 3 whose autistic child displayed the transmitted SNV in the *APH1A* gene. Family 4 whose autistic child displayed the *de novo* SNVs in the *APOA1* and *PHACTR3* genes. Family 15 whose autistic child displayed transmitted SNVs in the *PTK7, P2RX1* and *CTTNBP2* genes. Family 20 whose autistic child displayed the transmitted CNV within the *CHD8* gene, the transmitted SNV in the *SETD5* gene and *de novo* SNV in the *TMEM11* gene. Family 21 whose autistic child displayed the transmitted mutation in the *KTNAL2* gene. †de NOVO SNV, § transmitted SNV, * CNV

As mutations in ASD strongly associated genes are present in one of the unaffected parents, one can suggest that these mutations are not sufficient to generate an ASD phenotype. Interestingly, for *FEZF2* allele-linked mutations, 1 to 3 transmitted mutations in genes strongly associated with ASD are seen to coincide with the *FEZF2* mutation (Figure 5), indicating that a multiple hit hypothesis can be favored over the hypothesis of a single hit in a gene strongly associated with ASD and sufficient to cause disease.

### Spatiotemporal co-expression of mutated high-risk genes in *FEZF2* allele-linked and unlinked families

Characterization of the spatio-temporal pattern of expression of genes found mutated in our study can give important insights into disease mechanisms. Multiple studies demonstrated convergence of the highest-confidence non-syndromic ASD risk genes in the frontal lobe (i.e., prefrontal and primary motor cortex) in deep-layer projection neurons from 10 to 24 post-conception weeks (PCW), or approximately spanning the end of the early fetal period and the entire mid-fetal period (40–42).

We took advantage of t-distributed stochastic neighbor embedding methods by applying BrainScope (43) to explore high-dimensional data (dual t-SNE) across multiple brain regions and developmental periods from the BrainSpan consortium (44). With this approach, we were able to distinguish different clusters for the 65 ASD risk genes defined by Sanders et al 2015 suggesting that they display distinct spatiotemporal trajectories (**Figure 7A-B**). We identified a first group of genes (red) that clusters around *FEZF2* and a second group (black) that clusters around *KATNAL2. De novo* SNVs found either in *FEZF2* allele-linked and unlinked families segregated in these two different clusters.

**Figure 7.**
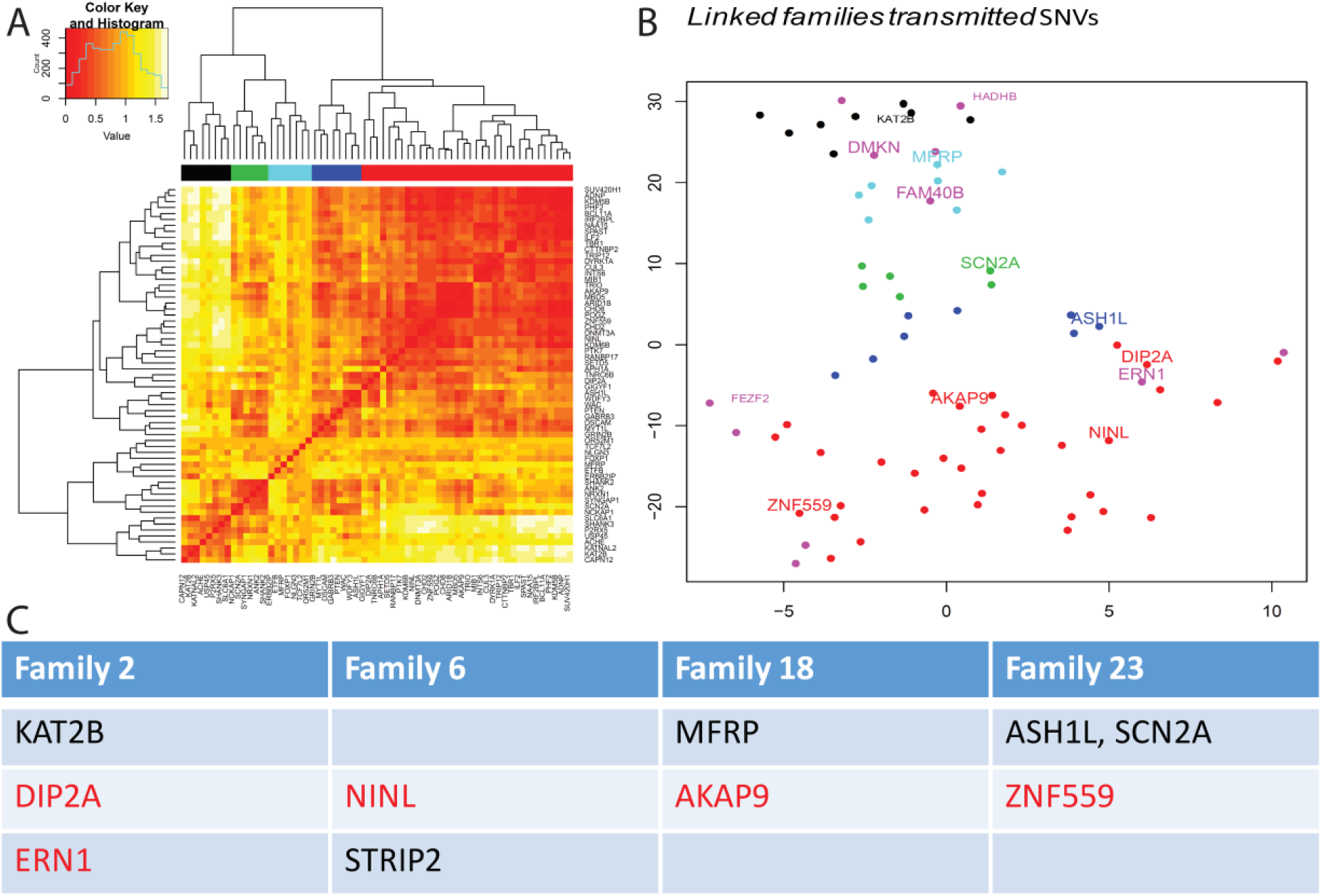
FEZF2 allele-linked families have a second hit from the 65 ASD risk genes defined by Sanders et al 2015 with spatio-temporal expression trajectories similar to that of *FEZF2*. A dual t-distributed stochastic neighbor embedding (t-SNE) method was used to analyze the 65 ASD high risk genes defined by Sanders et al., 2015, as well as the *de novo* and transmitted SNVs identified in our study. We used BrainScope (Huisman SMH et al., 2017 PMID: 28132031) to explore high-dimensional data (dual t-SNE) across developmental stages from the BrainSpan resource (Li et al., 2018 PMID: 30545854 and www.brainspan.org). (**A**) With this approach, we were able to distinguish different clusters for the 65 ASD risk genes defined by Sanders et al. 2015 suggesting that they display distinct spatiotemporal trajectories. (**B**) We identified a first group of genes (black) that clusters around KATNAL2 and a second group (red) that clusters around FEZF2. *De novo* SNVs found either in FEZF2 allele-linked and – unlinked families segregated in the different clusters. (**C**) Subset of the 65 ASD high-risk genes that were found to have mutations and *de novo* SNVs in the different FEZF2 allele-linked families. Genes that cluster around FEZF2 (red ensemble) are indicated in red, those that cluster around KATNAL2 (black ensemble) are indicated in black.

We focused our analysis on transmitted high-risk genes and *de novo* SNVs identified in *FEZF2* allele linked-families. In the red cluster, we identified *DIP2A* (Family 2), *NINL* (Family 6), AKAP9 (Family 18), and *ZNF559* (Family 23). In the black cluster, we identified *KAT2B* (Family 2), *MFRP* (Family 18), *ASH1L* and *SCN2A* (for Family 23); (**Figure 7C**). *De novo* SNVs in *ERN1* (Family 2) and *STRIP2* (Family 6) belong to the red and black ensembles, respectively (**Figure 7 C**).

Heatmap analysis of the spatiotemporal trajectories of genes displaying *de novo* mutations in the ‘red’ and ‘black’ clusters revealed contrasting patterns. The pattern for the red cluster is similar to that of *FEZF2* with spatiotemporal convergence in mid-fetal deep layer (Layer 5 and Layer 6) glutamatergic cortical pyramidal neurons (36) (**Supplementary Figure 2 A and C**). In contrast, the heatmap of the black ensemble is similar to that of *KATNAL2* with high levels of postnatal expression particularly in the striatum (**Supplementary Figure 2 B and D**).

Multiple sequence alignment for different proteins from phylogenetically distinct species indicated that the amino acid affected by the single nucleotide mutation was conserved through evolution, suggesting an important function for such amino acids (**Supplementary Figures 3-12**). Interestingly, *ZNF559* (Red ensemble, Family 23) is a primate-specific gene belonging to a cluster of KRAB zinc finger protein (KZFP) genes, involved in gene regulation and is not older than 100 million years (45) (**Supplementary Figure 13**). Regarding *SCN2A* (black ensemble, Family 23), mice with haploinsufficiency for the gene *Scn2a* displayed impaired spatial memory (46). A missense mutation, D12N, in *SNC2A* in the same domain as our A34T mutation (N terminal intra-cytoplasmic domain) appears to be able to generate an ASD phenotype in humans with functional defects of sodium channels (47). Furthermore, the product of *Scn2a*, NaV1.2 is critical for dendritic excitability and synaptic function in mature pyramidal neurons in addition to regulating early developmental axonal excitability (48). From transmitted CNVs, we identified a mutated CHD8 gene in *FEZF2* allele-unlinked family 20. This mutation involves the full deletion of a CHD8 exon that follows the helicase domain (**Supplementary Figure 14**). To further test the multiple-hit hypothesis in *FEZF2* allele-linked families 2, 6, 18 and 23, we investigated whether in all *FEZF2* allele-linked families at least one of those second hits was also expressed in *FEZF2*-expressing cells. For this analysis, we took advantage of single-cell and single-nuclei data from fetal and adult human frontal cortex (49, 50). In these datasets, *FEZF2* is expressed in neural progenitor cells (NPC) and excitatory neurons (L5 in adult). All families carried mutations in at least one gene that is expressed in the same developmental window as FEZF2 and are also co-expressed with FEZF2 in the same cell-type (**Figure 8**).

The layer distribution of these genes was further assessed using the 17 human cortex sections microarrays from (51) (**Figure 9**). The analysis of the relative expression of *FEZF2* across cortical sections confirmed its enrichment in layer 5. *AKAP9, ASH1L* and *SCN2A* were relatively more expressed in deeper sections of layer 5 but, contrary to *FEZF2*, none of them can be considered as significant layer specific markers in humans (51). *KAT2B* and *DIP2A* were enriched in L6 and described as L6 markers (51) despite differences in global RPKM (Reads Per Kilobase Million) expression levels (**Supplementary Figure 15**). *DIP2A* displays a higher number of reads in deep layer compared to upper layer cells in the adult frontal cortex (**Figure 8**). The layer differences observed in *KAT2B* might be contributed by glial cells (**Figure 8**). Independently of the degree of single-cell level expression overlap between *FEZF2* and putative second hits, none of the genes displayed the exact same cortical layer distribution of *FEZF2*.

**Figure 8.**
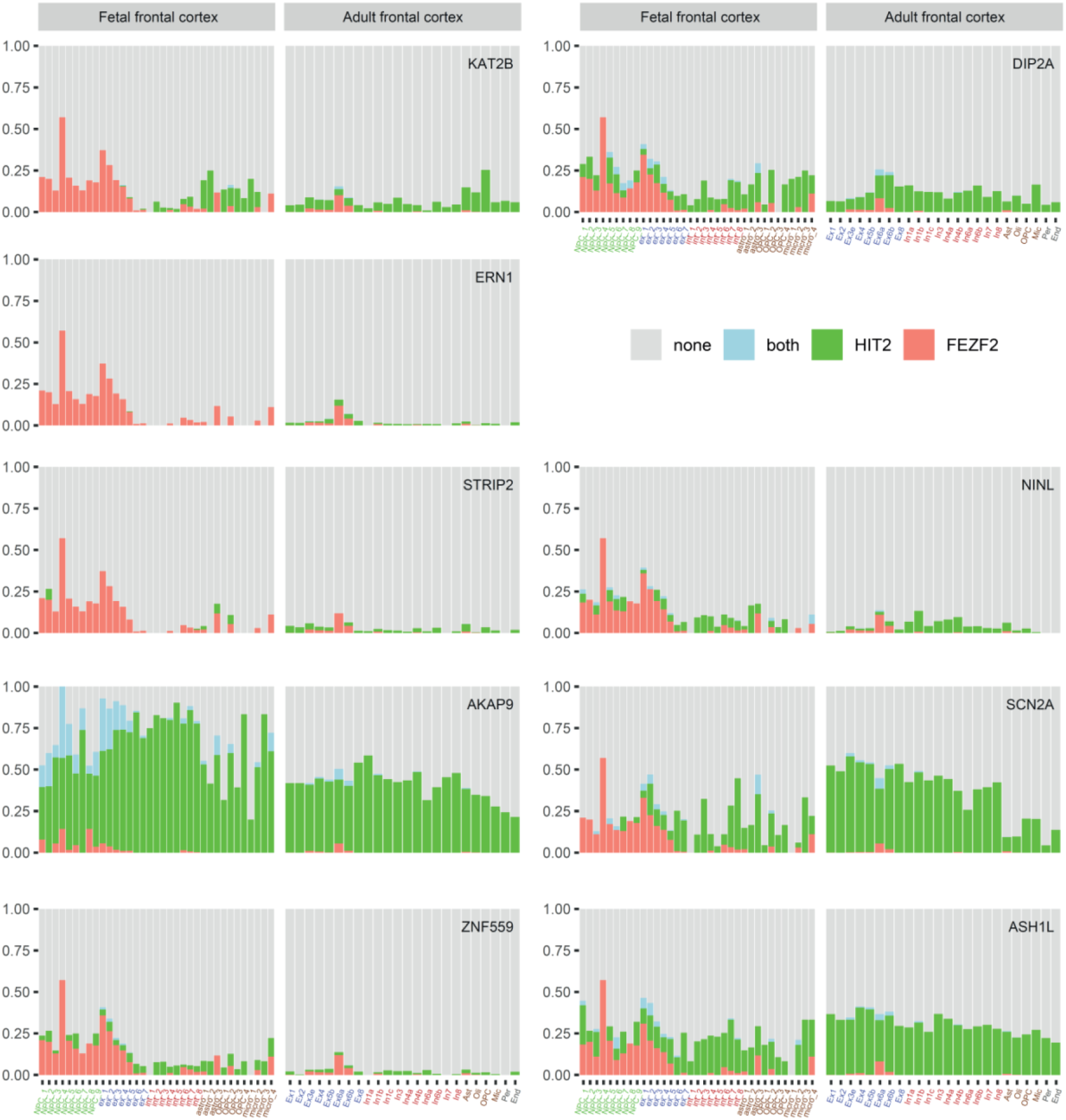
Co-expression of *FEZF2*, other mutated genes from the high-risk ensemble and genes displaying *de novo* SNVs in *FEZF2* allele-linked families. Barplots indicating the proportion *FEZF2*-expressing (red), “putative second hit”-expressing (green), and both *FEZF2*/second gene-expressing (blue) cells in frontal cortex single cell datasets derived from human fetal (Zhong et al. 2018) and adult (Lake et al. 2017) brains. High FEZF2 expression is found in progenitor cells and excitatory neurons in the fetal brain, and in cell cluster *Ex6a* in the adult brain, corresponding to a layer 5 subpopulation, as described in Lake et al. 2017.

**Figure 9.**
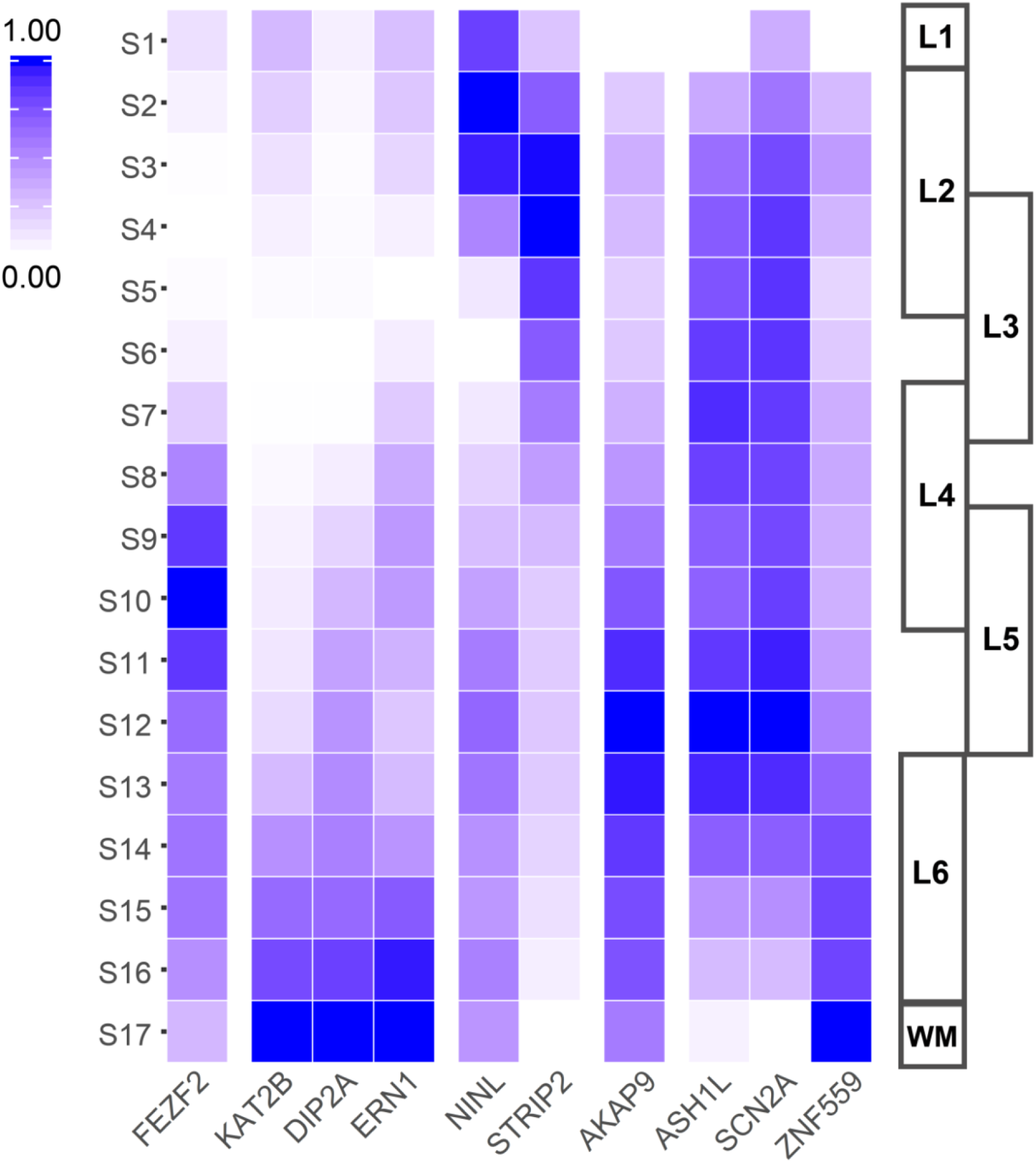
Heatmap showing relative gene expression levels of *FEZF2*, other mutated genes from the high-risk ensemble and *de novo* SNVs in *FEZF2* allele-linked families, across 17 prefrontal cortical sections of adult human brain. We used data from He et al. 2017, to create a layer expression profile for *FEZF2* and nine other ASD associated genes. The expression of *FEZF2* appears restricted to layer 5.

## Discussion

The genetic architecture of ASDs remains to be fully understood in spite of the identification of contributing rare and common variants (1, 2, 5). Our approach is based on the analysis of consanguineous families from Morocco whereas most other studies involve European and/or North American populations. The 1000 genomes project does not involve genomic data from Northern African countries. It is important to note the high level of CNVs in the Moroccan families analyzed in this study as compared to non-consangineous populations (21). Similar high level of CNVs have been reported in studies involving consanguineous families, such as in Lebanon (52).

Our aim was to test the hypothesis that a single damaging mutation in a limited set of high-risk genes, asproposed in previous studies (5, 6, 35), can be sufficient to cause ASD. Our data reveal a clear multi-hit organisation involving a homozygous *FEZF2* allele, a damaging *FEFZ2* mutation, mutations in ASD high-risk genes, and other gene variants. Each of these variants can contribute to the genesis of an ASD phenotype. The homozygous *FEZF2* allele may possibly impact on cis elements involving deregulation of the *FEZF2* gene. Such cis-regulatory structural variants associated with autism have been recently described (53). *FEZF2* is also part of a developing human brain gene co-expression network enriched for neurodevelopmental disorder risk genes that includes ASD high-risk genes such as *TBR1, CTTNBP2*, and *DSCAM* (44). Previously, genetic association has been found between *FEZF2* and ASD (54) and rare mutations in the *FEZF2* gene were identified in distinct ASD cohorts (55,56). The identification of a *FEFZ2* mutation reported here underscores the importance of *FEZF2* in ASD genetic architecture.

Considering our multi-hit model, further work will be needed to define the relative contribution for each mutation: the *FEZF2* mutation, one to three mutations in ASD high-risk genes, and one (or no) *de novo* SNVs identified in *FEZF2* allele-linked families (**Figure 7C**). For some ASD high-risk genes, such as *CDH8, SCN2A* and *SERD2*, their haploinsufficiency in mice is sufficient to induce autistic-like phenotypes as demonstrated for *CHD8* (57, 58), *SCN2A* (46), and *SETD5* (59). In our case, novel mouse models that can integrate the mutations reported here can be instrumental to dissect the contribution of each mutation to phenotypes relevant of ASD. Interestingly, in each FEZF2 allele-linked family we identified a mutation in a gene that is part of the FEZF2 ‘red ensemble’ and a mutation in a gene that is part of the *KATNAL2* ‘black ensemble’ (**Figure 5** and **Supplementary Figure 2**). Genes of the *FEZF2* ‘red ensemble’ that are expressed at midgestation in deep layers of the cortex may be involved in social interaction impairment as demonstrated for CDH8 (44). In contrast, genes of the *KATNAL2* ‘black ensemble’ that are highly expressed at postnatal stages in the striatum may be responsible for stereotypies (60, 61). How the mutations identified in *FEZF2* allele-linked families contribute to layer organisation of the human cortex can be studied using induced Pluripotent Stem Cells (iPSCs). These pluripotent stem cells show a remarkable ability to self-organize and differentiate *in vitro* in three-dimensional aggregates, known as organoids or organ spheroids, and to recapitulate aspects of human brain development and function (62, 63). Analysis of deep layer organization enables one to define the involvement of mutated genes discovered in patients of *FEZF2* allele-linked families.

Altogether, our results highlight the genetic architecture of ASD involving a founder mutation that can be combined with other variants to increase the risk of ASD. We propose *FEZF2* as a novel candidate to be sequenced in ASD, particularly in at-risk consanguineous families.

## Materials and Methods

### Patients and Assessments

Our data set consisted of 31 individuals from 10 consanguineous Moroccan ASD families recruited from the Child Psychiatry Center at the Arrazi Hospital (Salé –Morocco). Core inclusion criteria for ASD individuals were as follows: (1) clinical symptoms for more than 3 years, (2) a presumptive clinical diagnosis of ASD, (3) an expert clinical determination of ASD diagnosis based on DSM-IV criteria supported by the Autism Diagnosis Interview (ADI-R). All patients underwent clinical and molecular evaluation including fragile X testing and karyotype with normal results. Written informed consent was obtained from parents. Interviews of all participants were done using a protocol approved by the ethics committee at the Faculty of Medicine and Pharmacy in Rabat.

### Microarrays and copy number analysis

Genomic DNA was extracted from peripheral blood using a standard organic Phenol/Chloroform extraction. Genotyping with Affymetrix Genome Wide Human SNP Array 6.0 was performed using the protocol recommended by the manufacturer. In brief, a total of 2 × 250 ng of genomic DNA was used as starting material for each enzymatic digestion (Nsp and Sty) followed by linker-ligation of a common adaptor sequence. Standard PCR amplifies a predictable range of fragment sizes. The target was fragmented, hybridized to microarray, stained, and scanned. Quality controls were carried out. Affymetrix software Genotyping Console version 4.1 was used for genotype calling, quality control, and CNV identification. The genotyping analysis was performed using the Birdseed v2 algorithms. The samples were simultaneously genotyped with 90 Caucasian samples from the set of 270 HapMap individuals. Linkage analysis was performed using the Merlin software package.

Analysis of copy number and loss of heterozygosity were performed using the BRLMM-P-Plus algorithms. Each sample was compared to a reference file provided by Affymetrix and created from the 270 HapMap samples. CNV calls were generated via PennCNV.

### Selecting SNVs from Polyphen2 analysis

We used PolyPhen-2 that is a new development of the PolyPhen tool for annotating coding non synonymous SNPs ((Adzhubei et al., 2010). http://genetics.bwh.harvard.edu/pph2 Our criteria were (i) to have a possibly damaging in at least one of DIV and VAR parameters (pval <01) and (ii) to be a rare variant (frequency <0.01 in dbsnp38).

### PCR amplification and sequencing

PCR amplifications were performed with 50ng of genomic DNA using Platinum Taq DNA Polymerase (Invitrogen) and 10µM of each human FEZF2 primer. PCR products were purified and sequenced by GATC Biotech company.

### Assaying the function of the mutant 12bp deletion (d12bp) FEZF2 protein in a mouse model of corticospinal tract development

We made a construct with the 12bp deletion of FEZF2 able to generate a FEFZF2 protein with the RLPA deletion (see **Supplementary Table S4**). Both the wildtype and mutated human FEZF2 sequences were cloned into a mammalian expression vector, pCAGEN. Generation of Fezf2 Conditional Knockout Mice Fezf2^fl/fl^ has been described earlier (10). Briefly, E13.5 Fezf2^fl/fl^ mouse embryos were electroporated *in utero* with a combination of three expression constructs; pCAG-Cre to excise the endogenous mouse Fezf2 gene, pCALNL-GFP to visualize transfected cells, and either pCAG-FEZF2 to express the wildtype human FEZF2 protein or pCAG-d12bp-FEZF2 to express the mutant FEZF2 protein carrying the 12bp deletion. Electroporations were targeted to layer 5 of the cortex. The effect on corticospinal tract development was assessed at P0 by analysis of GFP-labelled projections in coronal brain sections (cortex and pons, respectively).

### Whole exome sequencing

We performed exome-sequencing with Illumina HISEQ using the manufacturer’s protocol. Quality control, alignment and calling were made with GATK suite. We used a Bayesian approach for filtering out false-positive calls. We used microarray data and exome-sequencing calls in order to establish two classes of calls: true positive (TP) and false-positive (FP). If the call in the microarray is equal to the call in the exome-sequencing we class it TP, otherwise we class it FP. We classed calls by the average coverage by site, i.e. we divide by 31 (the number of individuals) the total number of reads for a site, we separate heterozygous calls from homozygous calls. According to DP mean and states (heterozygous or homozygous), we constructed empirical distributions for GATK parameters and for TP and FP. We estimated a Bayes Factor (BF) for the call and we filtered out all calls with a log(BF)<3 and read depth<20. We used Sanger sequencing to perform validation of variants.

### Networks bioinformatics analyses

We used DAPPLE and WebGestalt suites. Disease Association Protein-Protein Link Evaluator (DAPPLE) looks for significant physical connectivity among proteins encoded for by genes in loci associated to disease according to protein-protein interactions reported in the literature (22). Interactions are extracted from the database “InWeb” that combines data from a variety of public PPI sources including MINT, BIND, IntAct and KEGG and defines high confidence interactions as those seen in multiple independent experiments. Connections can be direct (two proteins are physically linked to each other) and indirect (interaction is mediated by a common interactor). The non-randomness of the network and the significance of the interaction parameters are tested using a permutation method that compares the original network with thousands of networks created by randomly re-assigning the protein names while keeping the overall structure (size and number of interactions) of the original network. To complement the DAPPLE analysis, we used Over-Representation Analysis (ORA) from WebGestalt suite (31).

### t-distributed stochastic neighbor embedding methods

We took advantage of t-distributed stochastic neighbor embedding methods by applying BrainScope (43) to explore high-dimensional data (dual t-SNE) across multiple brain regions and developmental periods from the BrainSpan consortium (44).

### Gene co-expression analysis

Cell gene counts from fetal and adult human frontal cortex were obtained from Zhong et al. 2018 and Lake et al. 2017, respectively (49, 50). Counts were log-normalized using Seurat V3 (Stuart et al. 2019) and a scale factor of 10,000. In order to calculate the proportion of cells expressing or not expressing a given gene, a gene was considered expressed in one cell if its normalized value was greater than or equal to 1.

### Cortical layer expression

RPKM from the 17 cortical sections were obtained from He et al. 2017 (51). To represent relative expression across sections for each gene, we scaled RPKM values to be between 0 and 1 by subtracting the minimum and dividing by the maximum minus the minimum.

### Haplotype Network reconstruction

Network reconstruction of all carrier and non-carrier haplotypes was obtained using the median-joining algorithm implemented in NETWORK v.4.5.1 (Bandelt et al., 1999).

## ACKNOWLEDGMENTS

We are grateful to the families who participated in this study. We thank M Maad, S Amrani, F El Kabakbi, D Mehni, K Belmahjoub, A Sehnaji,, A Hachimi and also Collectif Autisme Maroc, Association Autisme 2005 (Meknès), Association Yahya pour Enfants Autistes (Tétouan), Association Idmaj Autiste (Casablanca), for help in awareness of families.

We thank Pr Fowzan S Alkuraya and Dr Hanan E Shamseddin, King Faisal Specialist Hospital and Research Center, Saudi Arabia King Saud University, Saudi Arabia King Fahad Medical City, Saudi Arabia for sharing data about FEZF2 mutations from Saudi Arabia patients cohorts.

## Legends of Supplementary Figures

**Supplementary Figure 1. Interactome network from products of genes found in *de novo* and transmitted CNVs for *FEZF2* allele-unlinked (A) and linked (B) families**.

Disease Association Protein-Protein Link Evaluator (DAPPLE) Network Derived from Genes hits found statistically significant networks linked to synapse for transmitted and *de novo* CNVs in *FEZF2* allele-unlinked families (A). In contrast, a significant DAPPLE Network Derived from Genes hit is found only for repertoire involving both *de novo* and transmitted CNVs in *FEZF2* allele-linked families (B).

**Supplementary Figure 2. Spatiotemporal expression of ASD high-risk genes from 65 genes list of Sanders et al. 2015**

We analysed spatiotemporal expression pattern trajectories of *FEZF2* (A) and *KATNAL2* (B) using the Brainscope resource (Huisman SMH et al., 2017 PMID: 28132031). Their pattern is similar to that of ‘red’ (C) and ‘black’ (D) ensembles, respectively, as defined in Figure 7.

**Supplementary Figure 3. Multiple sequence alignment of DIP2A proteins from different species**

We identified a mutation of the DIP2A protein (N553D) in Family n°2 linked to the *FEZF2* allele.

This mutation occurs in a region fully conserved from Xenopus to primates, including humans.

Multiple sequence alignment of DIP2A protein sequences from human, mouse, chimpanzee, rat, Macaca mulatta, Xenopus, Bos Taurus, Sumatran Orangutan, and Macaca fascicularis, respectively.

**Supplementary Figure 4. Multiple sequence alignment of NINL proteins from different species**

We identified a mutation of the NINL protein (E673Q) in the Family n°6 linked to the *FEZF2* allele.

This mutation occurs in a region conserved from rodents to primates, including humans.

Multiple sequence alignment of NINL protein sequences from human, mouse, chimpanzee, gorilla, Macaca nemestrina, Mandrillus leucophaeus, and Papio anubis, respectively.

**Supplementary Figure 5. Multiple sequence alignment of AKAP9 proteins from different species**

We identified a mutation of the AKAP9 protein (R1614Q) in the Family n°18 linked to the *FEZF2* allele.

This mutation occurs in a region fully conserved from cat to primates, including humans.

Multiple sequence alignment of AKAP9 protein sequences from human, mouse, rat, cat, Macaca mulatta, gorilla, Macaca fascicularis, pig, Papio anubis, Bonobo, and Macaca nemestrina, respectively.

**Supplementary Figure 6. Multiple sequence alignment of ZNF559 proteins from different species**

The *ZNF 559* gene is primate-specific. We identified a mutation of the ZNF559 protein (V318F) in Family n°23 linked to the *FEZF2* allele.

This mutation occurs in a region that includes a zinc finger motif (violet) and is fully conserved (yellow) in human and non-human primates (see here gorilla, Macaca mulatta, chimpanzee, Sumatran orangutan, Papio anubis, Macaca fascicularis, Black snub-nosed monkey, Peters’ Angolan colubus, Cercocebus, and Golden snub-nosed monkey, respectively).

**Supplementary Figure 7. Multiple sequence alignment of KAT2B proteins from different species**

We identified a mutation of the KAT2B protein (R653W) in Family n°2 linked to the *FEZF2* allele.

This mutation occurs in a region conserved from cat to primates, including humans.

Multiple sequence alignment of KAT2B protein sequences from Human, Macaca mulatta, cat, dog, pig, Papio Anubis, gorilla, Macaca Nemestrina, Sumatran orangutan, Giant panda, Bonobo, Cercopithecus sabeus, Macaca Fascicularis, and Bos taurus.

N-acetyltransferase domain from 503 to 651 is indicated in yellow

**Supplementary Figure 8. Multiple sequence alignment of MFRP proteins fromdifferent species**

We identified a mutation of the MFRP protein (L458F) in the Family n°18 linked to the *FEZF2* allele.

This mutation occurs in a region conserved from mouse to primates, including humans.

Multiple sequence alignment of MFRP protein sequences from Human, mouse, chimpanzee, rat, Bos taurus, Sumatran orangutan, Papio anubis, Macaca fascicularis, Bonobo, sheep, rabbit, Macaca nemestrina, pig, and Macaca mulatta.

The LDL-receptor class A 2 domain (420 – 455) and the Frizzled domain (461-579) are shown in yellow.

**Supplementary Figure 9. Multiple sequence alignment of ASH1L proteins from different species**

We identified a mutation of the ASH1L protein (L1136F) in the Family n°23 linked to the *FEZF2* allele.

This mutation occurs in a region conserved from mouse to primates, including humans.

Multiple sequence alignment of ASH1L protein sequences from Human, mouse, rat, dog, Bos taurus, horse, cat, Macaca mulatta, pig, Cercopithecus sabaeus, Giant Panda, Macaca fascicularis, Bonobo, and White-tufted ear marmoset.

**Supplementary Figure 10. Multiple sequence alignment of SCN2A proteins from different species**

We identified a mutation of the SCN2A protein (A34T) in Family n°23 linked to the *FEZF2* allele.

This mutation occurs in a region conserved from chick to primates, including humans.

Multiple sequence alignment of SCN2A protein sequences from Human, mouse, rat, dog, cat, chimpanzee, chick, pig, Macaca mulatta, rabbit, Giant panda, gorilla, Bonobo, and Macaca fascicularis.

The D12N mutation in the same region (N-terminal cytoplasmic tail) identified by Ben-Shalom et al., 2017 is indicated.

**Supplementary Figure 11. Multiple sequence alignment of ERN1 proteins from different species**

We identified a mutation of the ERN1 protein (S536L) in Family n°2 linked to the *FEZF2* allele. This mutation occurs in a region conserved from zebrafish to primates, including humans.

Multiple sequence alignment of ERN1 protein sequences from Human, mouse, Macaca mulatta, cat, rat, horse, chimpanzee, dog, Bos taurus, gorilla, Papio anubis, Macaca fascicularis, Bonobo, Macaca nemestrina, sheep, Giant panda, and Sumatran orangutan.

**Supplementary Figure 12. Multiple sequence alignment of STRIP2 proteins from different species**

We identified a mutation of the STRIP2 protein (I308V) in Family n°6 linked to the *FEZF2* allele.

This mutation occurs in a region conserved from zebrafish to primates, including humans.

Multiple sequence alignment of STRIP2 protein sequences from Human, mouse, dog, Macaca mulatta, cat, Bus Taurus, zebrafish, gorilla, chick, Giant panda, Sumatran orangutan, Papio anubis, horse, Macaca nemestrina, chimpanzee, and Bonobo.

**Supplementary Figure 13. The position of the *ZNF559* gene within the human genome and its primate-specific conservation**

Site of the *ZNF559* locus within the human genome (human HG38). Note that it is fully conserved in primates but not in other mammals, suggesting that *ZNF559* is a primate-specific innovation.

**Supplementary Figure 14. Localization of the deletion in *CHD8* gene identified in a FEZF2 allele-unlinked family**.

We identified a deletion in the *CHD8* gene in *FEZF2* allele-unlinked family 20. This mutation involves the full deletion of a *CHD8* exon that follows the helicase domain. (**A**). Localization of the 571bp deletion of the *CHD8* gene locus. (**B**) Deletion of 571bp indicating the *CDH8* exon involved and the phylogenetically conserved region.

**Supplementary Figure 15. Heatmap showing gene expression in RPKM across 17 prefrontal cortical sections**.

We used data from the adult human brain (He et al. 2017) to obtain profiles for *FEZF2* and ten other ASD associated genes.

## Tables

Supplementary Table S1: Clinical data

Supplementary Table S2: Linkage homozygosity

Supplementary Table S3: Haplotype counts

Supplementary Table S4: Sequences of FEZF2 and 12bpdel FEZF2 constructs

Supplementary Table S5: *De novo* and transmitted CNVs

Supplementary Table S6: *De novo* SNVs

Supplementary Table S7: List of CNVs found in more than one family

Supplementary Table S8: Transmitted SNVs for linked and unlinked families

## Methods references

Adzhubei IA, Schmidt S, Peshkin L, Ramensky VE, Gerasimova A, Bork P, Kondrashov AS, Sunyaev SR. (2010) Nat Methods 7(4):248–249.

Stuart T, Butler A, Hoffman P, Hafemeister C, Papalexi E, Mauck WM 3rd, Hao Y, Stoeckius M, Smibert P, Satija R (2019). Comprehensive Integration of Single-Cell Data. Cell. 177(7):1888–1902.e21.

Bandelt, H.J., Forster, P. and Rohl, A. (1999) Median-joining networks for inferring intraspecific phylogenies. Mol. Biol. Evol., 16, 37–48.

## REFERENCES

1. Vorstman JAS, et al. (2017) Autism genetics: opportunities and challenges for clinical translation. Nat Rev Genet 18(6):362–376.

2. Chaste P, Roeder K, Devlin B (2017) The Yin and Yang of Autism Genetics: How Rare De Novo and Common Variations Affect Liability. Annu Rev Genomics Hum Genet

3. Lord C, Elsabbagh M, Baird G, Veenstra-Vanderweele J (2018) Autism spectrum disorder. Lancet 392(10146):508–520.

4. Lyall K, et al. (2017) The Changing Epidemiology of Autism Spectrum Disorders. Annu Rev Public Health 38:81–102.

5. Sanders SJ, et al. (2015) Insights into Autism Spectrum Disorder Genomic Architecture and Biology from 71 Risk Loci. Neuron 87(6):1215–1233.

6. Willsey AJ, et al. (2018) The Psychiatric Cell Map Initiative: A Convergent Systems Biological Approach to Illuminating Key Molecular Pathways in Neuropsychiatric Disorders. Cell 174(3):505–520.

7. Sebat J, et al. (2007) Strong association of de novo copy number mutations with autism. Science 316(5823):445–449.

8. Morrow EM, et al. (2008) Identifying autism loci and genes by tracing recent shared ancestry. Science 321(5886):218–223.

9. Gamsiz ED, et al. (2013) Intellectual disability is associated with increased runs of homozygosity in simplex autism. Am J Hum Genet 93(1):103–109.

10. Chen B, Schaevitz LR, McConnell SK (2005) Fezl regulates the differentiation and axon targeting of layer 5 subcortical projection neurons in cerebral cortex. Proc Natl Acad Sci USA 102(47):17184–17189.

11. Molyneaux BJ, Arlotta P, Hirata T, Hibi M, Macklis JD, (2005) Fezl is required for the birth and specification of corticospinal motor neurons.Neuron. 47(6):817–31.

12. Chen JG, Rasin MR, Kwan KY, Sestan N. (2005). Zfp312 is required for subcortical axonal projections and dendritic morphology of deep-layer pyramidal neurons of the cerebral cortex. Proc Natl Acad Sci U S A.102(49):17792–7.

13. Anazi S, et al. (2017) Clinical genomics expands the morbid genome of intellectual disability and offers a high diagnostic yield. Mol Psychiatry 22(4):615–624.

14. Lesage S, et al. (2010). Parkinson’s disease-related LRRK2 G2019S mutation results from independent mutational events in human. Hum Mol Genet. 19(10):1998–2004

15. Han W, et al. (2011) TBR1 directly represses Fezf2 to control the laminar origin and development of the corticospinal tract. Proc Natl Acad Sci USA 108(7):3041–3046.

16. McKenna WL, et al. (2011). Tbr1 and Fezf2 regulate alternate corticofugal neuronal identities during neocortical development. J Neurosci. 31(2):549–64.

17. Srinivasan K, et al. (2012). A network of genetic repression and derepression specifies projection fates in the developing neocortex. Proc Natl Acad Sci U S A. 2012 Nov 20;109(47):19071–8. doi: 10.1073/pnas.1216793109.

18. Fazel Darbandi S, et al. (2018) Neonatal Tbr1 Dosage Controls Cortical Layer 6 Connectivity. Neuron 100(4):831–845.e7.

19. Shim S, Kwan KY, Li M, Lefebvre V, Sestan N (2012) Cis-regulatory control of corticospinal system development and evolution. Nature 486(7401):74–79.

20. Gu Z, et al. (2017) Control of species-dependent cortico-motoneuronal connections underlying manual dexterity. Science 357(6349):400–404.

21. Leppa VM, et al. (2016) Rare Inherited and De Novo CNVs Reveal Complex Contributions to ASD Risk in Multiplex Families. Am J Hum Genet 99(3):540–554.

22. Rossin EJ, et al. (2011) Proteins encoded in genomic regions associated with immune-mediated disease physically interact and suggest underlying biology. PLoS Genet 7(1):e1001273.

23. Lundby A, et al. (2014) Annotation of loci from genome-wide association studies using tissue-specific quantitative interaction proteomics. Nat Methods 11(8):868–874.

24. Südhof TC (2012) The presynaptic active zone. Neuron 75(1):11–25.

25. Südhof TC (2018) Towards an Understanding of Synapse Formation. Neuron 100(2):276–293.

26. Xu J, et al. (2017) Complete Disruption of the Kainate Receptor Gene Family Results in Corticostriatal Dysfunction in Mice. Cell Rep 18(8):1848–1857.

27. Nelson SB, Valakh V (2015) Excitatory/Inhibitory Balance and Circuit Homeostasis in Autism Spectrum Disorders. Neuron 87(4):684–698.

28. Nguyen Ba-Charvet KT, et al. (1999) Slit2-Mediated chemorepulsion and collapse of developing forebrain axons. Neuron 22(3):463–473.

29. Leshchyns’ka I, Sytnyk V (2016) Reciprocal Interactions between Cell Adhesion Molecules of the Immunoglobulin Superfamily and the Cytoskeleton in Neurons. Front Cell Dev Biol 4:9.

30. Leblond CS, et al. (2014) Meta-analysis of SHANK Mutations in Autism Spectrum Disorders: a gradient of severity in cognitive impairments. PLoS Genet 10(9):e1004580.

31. Wang J, Vasaikar S, Shi Z, Greer M, Zhang B (2017) WebGestalt 2017: a more comprehensive, powerful, flexible and interactive gene set enrichment analysis toolkit. Nucleic Acids Res 45(W1):W130–W137.

32. Gilissen C, et al. (2014) Genome sequencing identifies major causes of severe intellectual disability. Nature 511(7509):344–347.

33. Iossifov I, et al. (2014) The contribution of de novo coding mutations to autism spectrum disorder. Nature 515(7526):216–221.

34. De Rubeis S, et al. (2014) Synaptic, transcriptional and chromatin genes disrupted in autism. Nature 515(7526):209–215.

35. Sestan N, State MW (2018) Lost in Translation: Traversing the Complex Path from Genomics to Therapeutics in Autism Spectrum Disorder. Neuron 100(2):406–423.

36. Chen WV, Maniatis T (2013) Clustered protocadherins. Development 140(16):3297–3302.

37. Mountoufaris G, Canzio D, Nwakeze CL, Chen WV, Maniatis T (2018) Writing, Reading, and Translating the Clustered Protocadherin Cell Surface Recognition Code for Neural Circuit Assembly. Annu Rev Cell Dev Biol 34:471–493.

38. Rozenkrantz L, et al. (2015) A Mechanistic Link between Olfaction and Autism Spectrum Disorder. Curr Biol 25(14):1904–1910.

39. Endevelt-Shapira Y, et al. (2018) Altered responses to social chemosignals in autism spectrum disorder. Nat Neurosci 21(1):111–119.

40. Willsey AJ, et al. (2013) Coexpression networks implicate human midfetal deep cortical projection neurons in the pathogenesis of autism. Cell 155(5):997–1007.

41. Pinto D, et al. (2014) Convergence of genes and cellular pathways dysregulated in autism spectrum disorders. Am J Hum Genet 94(5):677–694.

42. Uddin M, et al. (2014) Brain-expressed exons under purifying selection are enriched for de novo mutations in autism spectrum disorder. Nat Genet 46(7):742–747.

43. Huisman SMH, et al. (2017) BrainScope: interactive visual exploration of the spatial and temporal human brain transcriptome. Nucleic Acids Res 45(10):e83.

44. Li M, et al. (2018) Integrative functional genomic analysis of human brain development and neuropsychiatric risks. Science 362(6420).

45. Pontis J, et al. (2019). Hominoid-Specific Transposable Elements and KZFPs Facilitate Human Embryonic Genome Activation and Control Transcription in Naive Human ESCs. Cell Stem Cell. 24(5):724–735.

46. Middleton SJ, et al. (2018) Altered hippocampal replay is associated with memory impairment in mice heterozygous for the Scn2a gene. Nat Neurosci 21(7):996–1003.

47. Ben-Shalom R, et al. (2017). Opposing Effects on NaV1.2 Function Underlie Differences Between SCN2A Variants Observed in Individuals With Autism Spectrum Disorder or Infantile Seizures. Biol Psychiatry. 82(3):224–232.

48. Spratt PWE, et al. (2019).The Autism-Associated Gene Scn2a Contributes to Dendritic Excitability and Synaptic Function in the Prefrontal Cortex. Neuron. 2 pii: S0896-6273(19)30490–8.

49. Zhong S, et al. (2018). A single-cell RNA-seq survey of the developmental landscape of the human prefrontal cortex. Nature. 555(7697):524–528.

50. Lake BB, et al. (2018). Integrative single-cell analysis of transcriptional and epigenetic states in the human adult brain.Nat Biotechnol. 36(1):70–80.

51. He Z, et al. (2017). Comprehensive transcriptome analysis of neocortical layers in humans, chimpanzees and macaques. Nat Neurosci.20(6):886–895.

52. Soueid J, et al. (2016) RYR2, PTDSS1 and AREG genes are implicated in a Lebanese population-based study of copy number variation in autism. Sci Rep 6:19088.

53. Brandler WM, et al. (2018) Paternally inherited cis-regulatory structural variants are associated with autism. Science 360(6386):327–331.

54. Wang K et al. (2009) Common genetic variants on 5p14.1 associate with autism spectrum disorders. Nature. 459(7246):528–33.

55. Sanders SJ et al. (2012). De novo mutations revealed by whole-exome sequencing are strongly associated with autism. Nature.485(7397):237–41.

56. Feliciano P et al. (2019). Exome sequencing of 457 autism families recruited online provides evidence for autism risk genes. NPJ Genom Med. Aug 23;4:19

57. Katayama Y, et al. (2016) CHD8 haploinsufficiency results in autistic-like phenotypes in mice. Nature 537(7622):675–679.

58. Gompers AL, et al. (2017) Germline Chd8 haploinsufficiency alters brain development in mouse. Nat Neurosci 20(8):1062–1073.

59. Deliu E, et al. (2018) Haploinsufficiency of the intellectual disability gene SETD5 disturbs developmental gene expression and cognition. Nat Neurosci 21(12):1717–1727.

60. Burguière E, Monteiro P, Feng G, Graybiel AM. (2013). Optogenetic stimulation of lateral orbitofronto-striatal pathway suppresses compulsive behaviors.Science. 340(6137):1243–6.

61. Kaiser T, Feng G.(2015) Modeling psychiatric disorders for developing effective treatments. Nat Med. 21(9):979–88.

62. Amin ND, Paşca SP. (2018). Building Models of Brain Disorders with Three-Dimensional Organoids.Neuron 100(2):389–405.

63. Yoon SJ et al.(2019). Reliability of human cortical organoid generation.Nat Methods. 16(1):75–78.

