## Supplementary Figures for "Multi-hit autism genomic architecture evidenced from consanguineous families with involvement of FEZF2 and mutations in high-risk genes"

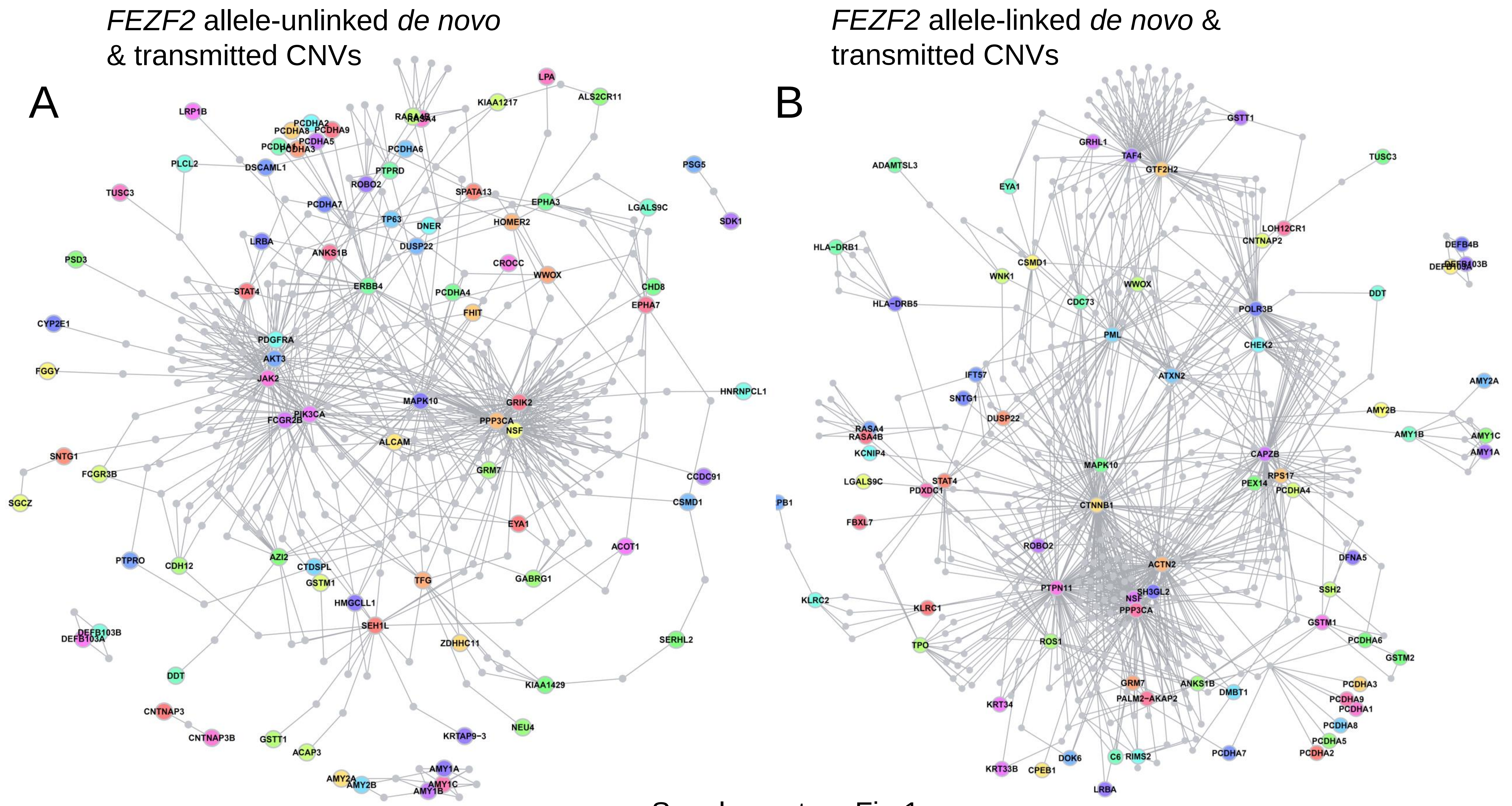


Supplementary Fig.1

**Supplementary Figure 1. Interactome network from products of genes found in *de novo* and transmitted CNVs for *FEZF2* allele-unlinked (A) and linked (B) families**.

Disease Association Protein-Protein Link Evaluator (DAPPLE) Network Derived from Genes hits found statistically significant networks linked to synapse for transmitted and de novo CNVs in *FEZF2* allele-unlinked families (A). In contrast, a significant DAPPLE Network Derived from Genes hit is found only for repertoire involving both de novo and transmitted CNVs in *FEZF2* allele-linked families (B).

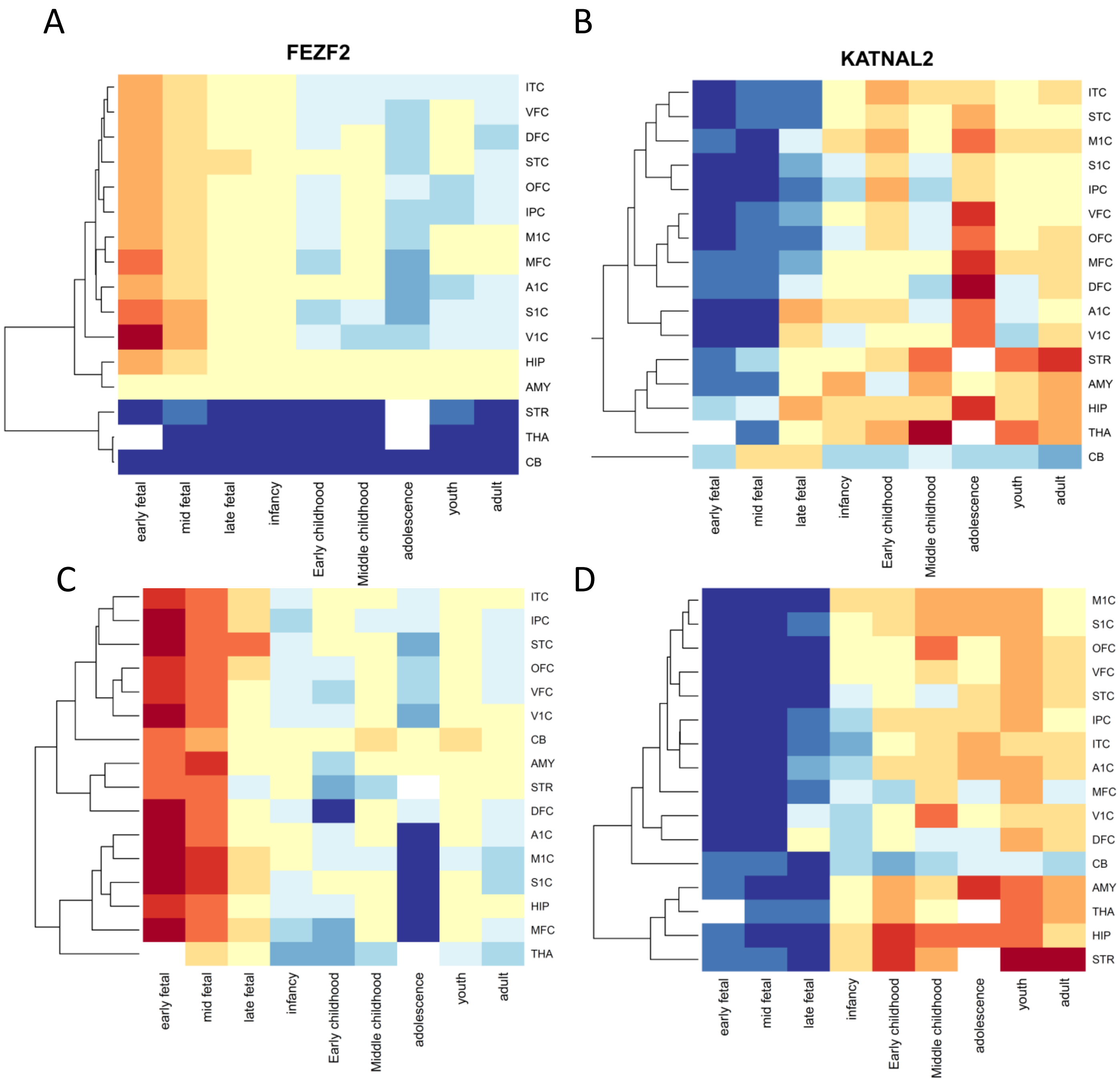


Supplementary Figure 2

**Supplementary Figure 2. Spatiotemporal expression of ASD high-risk genes from 65 genes list of Sanders et al. 2015**

We analysed spatiotemporal trajectories pattern of *FEZF2* (A) and *KATNAL2* (B) using Brainscope resource (Huisman SMH et al., 2017 PMID: 28132031). Their pattern is similar to that of ‘red’ (C) and ‘black’ (D) ensembles, respectively as defined in Figure 7.

### DIP2A protein N553D

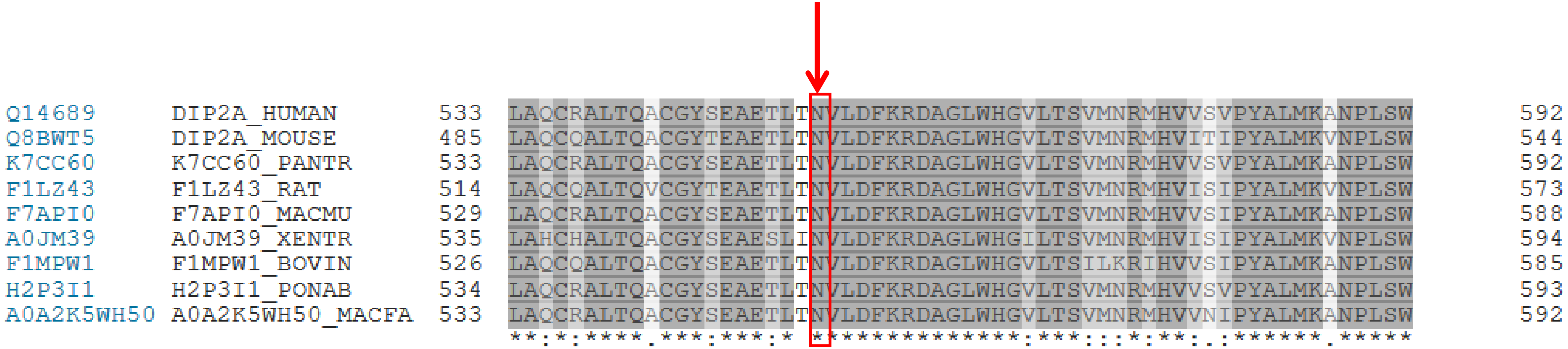


Supplementary Fig.3

**Supplementary Figure 3. Multiple sequence alignment of DIP2A proteins for different species**

We identified a mutation of DIP2A protein (N553D) in the family n°2 linked to *FEZF2* allele.

This mutation occurs in a region fully conserved from Xenopus to primates including humans.

Multiple sequence alignment of DIP2A protein sequences from human, mouse, chimpanzee, rat, Macaca mulatta, Xenopus, Bos Taurus, Sumatran Orangutan and Macaca fascicularis, respectively.

### NINL protein E673Q

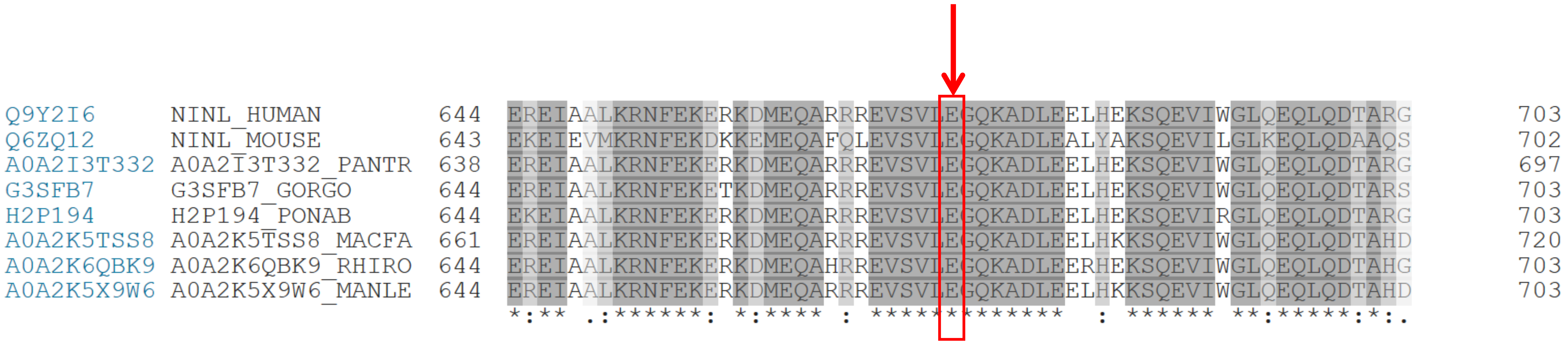


Supplementary Fig.4

**Supplementary Figure 4. Multiple sequence alignment of NINL proteins for different species**

We identified a mutation of NINL protein (E673Q) in the family n°6 linked to *FEZF2* allele.

This mutation occurs in a region conserved from rodents to primates including humans.

Multiple sequence alignment of NINL protein sequences from human, mouse, chimpanzee, gorilla, Macaca nemestrina, Mandrillus leucophaeus and Papio anubis, respectively.

AKAP9 protein R1614Q

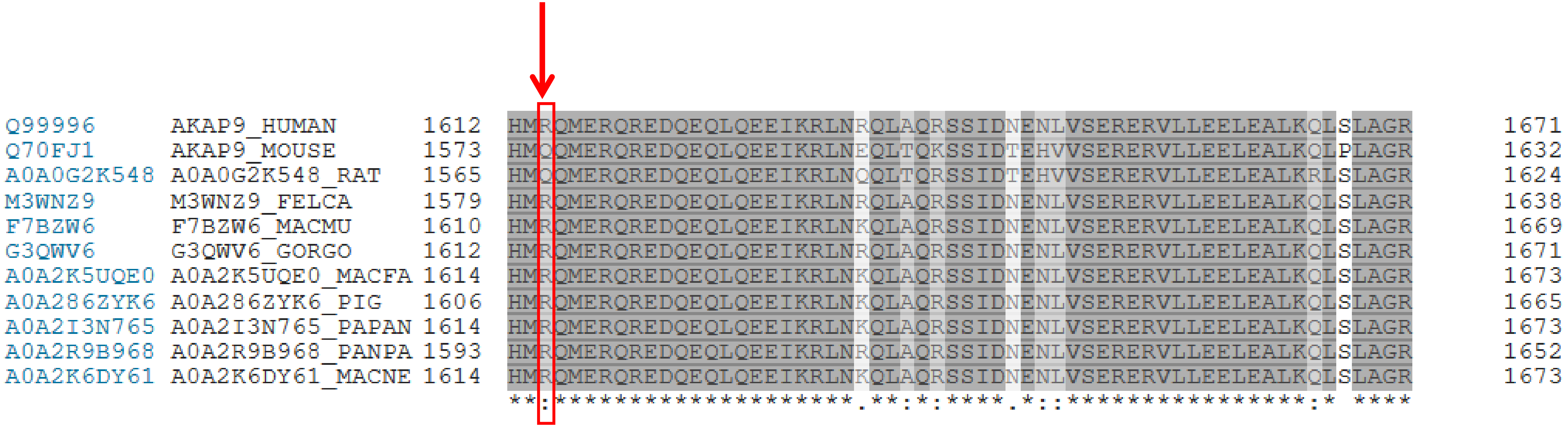


Supplementary Fig.5

**Supplementary Figure 5. Multiple sequence alignment of AKAP9 proteins for different species**

We identified a mutation of AKAP9 protein (R1614Q) in the family n°18 linked to *FEZF2* allele.

This mutation occurs in a region fully conserved from cat to primates including humans.

Multiple sequence alignment of AKAP9 protein sequences from human, mouse, rat, cat, Macaca mulatta, gorilla, Macaca fascicularis, pig, Papio anubis, Bonobo and Macaca nemestrina, respectively.

### ZNF559 protein V318F

```
Q9BR84      ZN559_HUMAN       296  HYVCNECGKEFTCFSKLNIHIRVHTGEKPYECNKCGKAFTDSSGLIKHRRTH----TGEK  351
G3RAY3      G3RAY3_GORGO      360  HYVCNECRKEFTCFSKLNIHIRVHTGEKPYECNKCGKAFTDSSGLIKHRRTH----TGEK  415
A0A1D5Q255  A0A1D5Q255_MACMU  294  HYVCNECGKEFTCFSKLNIHIRVHTGEKPYECNKCGKAFTDSSGLIKHRRTH----TGEK  349
H2R6D9      H2R6D9_PANTR      309  HYVCNECGKEFTCFSKLNIHIRVHTGEKPYECNKCGKAFTDSSGLIKHRRTH----TGEK  364
H2NXG3      H2NXG3_PONAB      296  HYVCNECGKEFTCFSKLNIHIRVHTGEKPYECNKCGKTFTDSSGLIKHRRTHCVLFTGEK  355
A0A0A0MX59  A0A0A0MX59_PAPAN  346  HYVCNECGKEFTCFSKLNIHIRVHTGEKPYECNKCGKAFTDSSGLIKHRRTH----TGEK  401
A0A2K5V758  A0A2K5V758_MACFA  356  HYVCNECGKEFTCFSKLNIHIRVHTGEKPYECNKCGKAFTDSSGLIKHRRTH----TGEK  411
A0A2K6LUH1  A0A2K6LUH1_RHIBE  294  HYVCNECGKEFTCFSKLNIHIRVHTGEKPYECNKCGKAFTDSSGLVKHRRTH----TGEK  349
A0A2K5J159  A0A2K5J159_COLAP  294  HYVCNECGKEFTCFSKLNIHIRVHTGEKPYECNKCGKAFTDSSGLIKHRRTH----TGEK  349
A0A2K5NCW1  A0A2K5NCW1_CERAT  322  HYVCNECGKEFTCFSKLSIHIRVHTGEKPYECNKCGKAFTDSSGLIKHRRTH----TGEK  377
A0A2K6NUC1  A0A2K6NUC1_RHIRO  312  HYVCNECGKEFTCFSKLNIHIRVHTGEKPYECNKCGKAFTDSSGLIKHRRTH----TGEK  367
                                   ******* *********.*************************:*******:******    ****
```

Supplementary Fig.6

**Supplementary Figure 6. Multiple sequence aligment of ZNF559 proteins for different species**

*ZNF 559* gene is primate-specific. We identified a mutation of ZNF559 protein (V318F) in the family n°23 linked to *FEZF2* allele.

This mutation occurs in a region fully conserved (yellow) that includes a zinc finger motif (violet) in human and non-human primates (see here gorilla, Macaca mulatta, chimpanzee, Sumatran orangutan, Papio anubis, Macaca fascicularis, Black snub-nosed monkey, Peters' Angolan colubus, Cercocebus and Golden snub-nosed monkey, respectively).

### KAT2B R653W

| | | | | |
|---|---|---|---|---|
| Q92831 | KAT2B_HUMAN | 647 | GCELNPRIPYTEFSVIIKKQKEIIKKLIERKQAQIRKVYPGLSCFKDGVRQIPIESIPGI | 706 |
| F6W9N2 | F6W9N2_MACMU | 624 | GCELNPRIPYTEFSVIIKKQKEIIKKLIERKQAQIRKVYPGLSCFKDGVRQIPIESIPGI | 683 |
| M3W281 | M3W281_FELCA | 649 | GCELNPRIPYTEFSVIIKKQKEIIKKLIERKQAQIRKVYPGLSCFKDGVRQIPIESIPGI | 708 |
| F1PN31 | F1PN31_CANLF | 555 | GCELNPRIPYTEFSVIIKKQKEIIKKLIERKQAQIRKVYPGLSCFKDGVRQIPIESIPGI | 614 |
| I3LRW1 | I3LRW1_PIG | 627 | GCELNPRIPYTEFSVIIKKQKEIIKKLIERKQAQIRKVYPGLACFKDGVRQIPIESIPGI | 686 |
| A0A096NY03 | A0A096NY03_PAPAN | 647 | GCELNPRIPYTEFSVIIKKQKEIIKKLIERKQAQIRKVYPGLSCFKDGVRQIPIESIPGI | 706 |
| G3RWA3 | G3RWA3_GORGO | 635 | GCELNPRIPYTEFSVIIKKQKEIIKKLIERKQAQIRKVYPGLSCFKDGVRQIPIESIPGI | 694 |
| A0A2K6B5H3 | A0A2K6B5H3_MACNE | 615 | GCELNPRIPYTEFSVIIKKQKEIIKKLIERKQAQIRKVYPGLSCFKDGVRQIPIESIPGI | 674 |
| H2PBC4 | H2PBC4_PONAB | 570 | GCELNPRIPYTEFSVIIKKQKEIIKKLIERKQAQIRKVYPGLSCFKDGVRQIPIESIPGI | 629 |
| G1L279 | G1L279_AILME | 569 | GCELNPRIPYTEFSVIIKKQKEIIKKLIERKQAQIRKVYPGLSCFKDGVRQIPIESIPGI | 628 |
| A0A2R9BZT9 | A0A2R9BZT9_PANPA | 490 | GCELNPRIPYTEFSVIIKKQKEIIKKLIERKQAQIRKVYPGLSCFKDGVRQIPIESIPGI | 549 |
| A0A0D9RBQ9 | A0A0D9RBQ9_CHLSB | 490 | GCELNPRIPYTEFSVIIKKQKEIIKKLIERKQAQIRKVYPGLSCFKDGVRQIPIESIPGI | 549 |
| A0A2K5VBM7 | A0A2K5VBM7_MACFA | 546 | GCELNPRIPYTEFSVIIKKQKEIIKKLIERKQAQIRKVYPGLSCFKDGVRQIPIESIPGI | 605 |
| A0A3Q1MLT2 | A0A3Q1MLT2_BOVIN | 628 | GCELNPRIPYTEFSVIIKKQKEIIKKLIERKQAQIRKVYPGLSCFKDGVRQIPIESIPGI | 687 |
| | | | ******************************************:***************** | |

Supplementary Fig.7

**Supplementary Figure 7. Multiple sequence alignment of KAT2B proteins for different species**

We identified a mutation of KAT2B (R653W) in the family n°2 linked to *FEZF2* allele.
This mutation occurs in a region conserved from cat to primates including humans.

Multiple sequence alignment of KAT2B protein sequences from Human, Macaca mulatta, cat, dog, pig, Papio Anubis, gorilla, Macaca Nemestrina, Sumatran orangutan, Giant panda, Bonobo, Cercopithecus sabeus, Macaca Fascicularis and Bos taurus.

N-acetyltransferase domain from 503 to 651 is indicated in yellow

### MFRP L458F

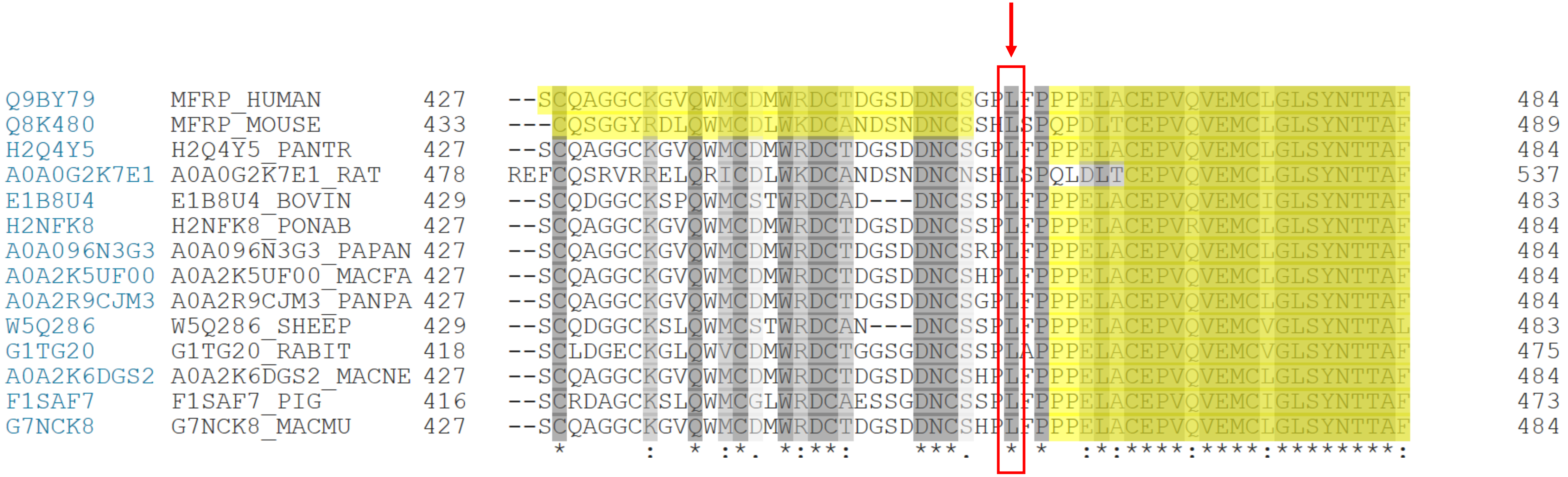


Supplementary Fig.8

**Supplementary Figure 8. Multiple sequence alignment of MFRP proteins for different species**

We identified a mutation of MFRP (L458F) in the family n°18 linked to *FEZF2* allele.
This mutation occurs in a region conserved from mouse to primates including humans.

Multiple sequence alignment of MFRP protein sequences from Human, mouse, chimpanzee, rat, Bos taurus, Sumatran orangutan, Papio anubis, Macaca fascicularis, Bonobo, sheep, rabbit, Macaca nemestrina, pig and Macaca mulatta.

LDL-receptor class A 2 domain (420 – 455) and Frizzled domain (461-579) are in yellow.

### ASH1L L1136F

```
Q9NR48       ASH1L_HUMAN        1079  QAAGSALGQILPPLLPSSASSSEILPSPICSQSSGTSGGQSPVSSDAGFVEPSSVPYLHL  1138
Q99MY8       ASH1L_MOUSE        1077  QAAGSALGQILPPLLPSPASSSEILPSPICSQSSGTSGGQSPVSSDAGFVEPSSVPYLHV  1136
D3ZKH4       D3ZKH4_RAT         1077  QAAGSALGQILPPLLPSPASSSEILPSPICSQSSGTSGGQSPVSSDAGFVEPSSVPYLHV  1136
E2RS85       E2RS85_CANLF       1079  QAAGSALGQILPPLLPSSASSSEILPSPVCSQSSGTSGGQSPVSSDAGFVEPSSVPYLHL  1138
E1BGA4       E1BGA4_BOVIN       1079  QAAGSALGQILPPLLPSSASSSEILPSPICSQSSGTSGGQSPVSSDAGFVEPSSVPYLHL  1138
F7A5F1       F7A5F1_HORSE       1079  QAAGSALGQILPPLLPSSASSSEILPSPICSQSSGTSGGQSPVSSDAGFVEPSSVPYLHL  1138
A0A2I2UW79   A0A2I2UW79_FELCA   1078  QAAGSALGQILPPLLPSSASGSEILPSPVCSQSSGTSGGQSPVSSDAGFVEPSSVPYLHL  1137
H9FX47       H9FX47_MACMU       1078  QATGSALGQILPPLLPSSASSSEILPSPICSQSSGTSGGQSPVSSDAGFVEPSSVPYLHL  1137
F1RLM3       F1RLM3_PIG         1073  QATGSALGPILPPLLPSSASSSEILPSPICSQSSGTSGGQSPVSSDAGFVEPSSVPYLHL  1132
A0A0D9S570   A0A0D9S570_CHLSB   1079  QATGSALGQILPPLLPSSASSSEILPSPICSQSSGTSGGQSPVSSDAGFVEPSSVPYLHL  1138
G1KZK0       G1KZK0_AILME       1079  QAAGSALGQILPSLLPSSASSSEILPSPICSQSSGTSGGQSPVSSDAGFVEPSSVPYLHL  1138
A0A2K5UCW9   A0A2K5UCW9_MACFA   1078  QATGSALGQILPPLLPSSASSSEILPSPICSQSSGTSGGQSPVSSDAGFVEPSSVPYLHL  1137
A0A2R9ASA3   A0A2R9ASA3_PANPA   1055  QAAGSALGQILPPLLPSSAGSSEILPSPICSQSSGTSGGQSPVSSDAGFVEPSSVPYLHL  1114
F7HAH4       F7HAH4_CALJA       1080  QATGSALGQILPPLLPSSASSSEILPSPICSQSSGTSGGQSPVSSDAGFVEPSSVPYLHL  1139
                                      **:***** *** **** *..*******:******************************:
```

Human, mouse, rat, dog, Bos taurus, horse, cat, Macaca mulatta, pig, Cercopithecus sabaeus, Giant Panda, Macaca fascicularis, Bonobo and White-tufted ear marmoset.

Supplementary Fig.9

**Supplementary Figure 9. Multiple sequence alignment of ASH1L proteins for different species**

We identified a mutation of ASH1L (L1136F) in the family n°23 linked to *FEZF2* allele.

This mutation occurs in a region conserved from mouse to primates including humans.

Multiple sequence alignment of ASH1L protein sequences from Human, mouse, rat, dog, Bos taurus, horse, cat, Macaca mulatta, pig, Cercopithecus sabaeus, Giant Panda, Macaca fascicularis, Bonobo and White-tufted ear marmoset.

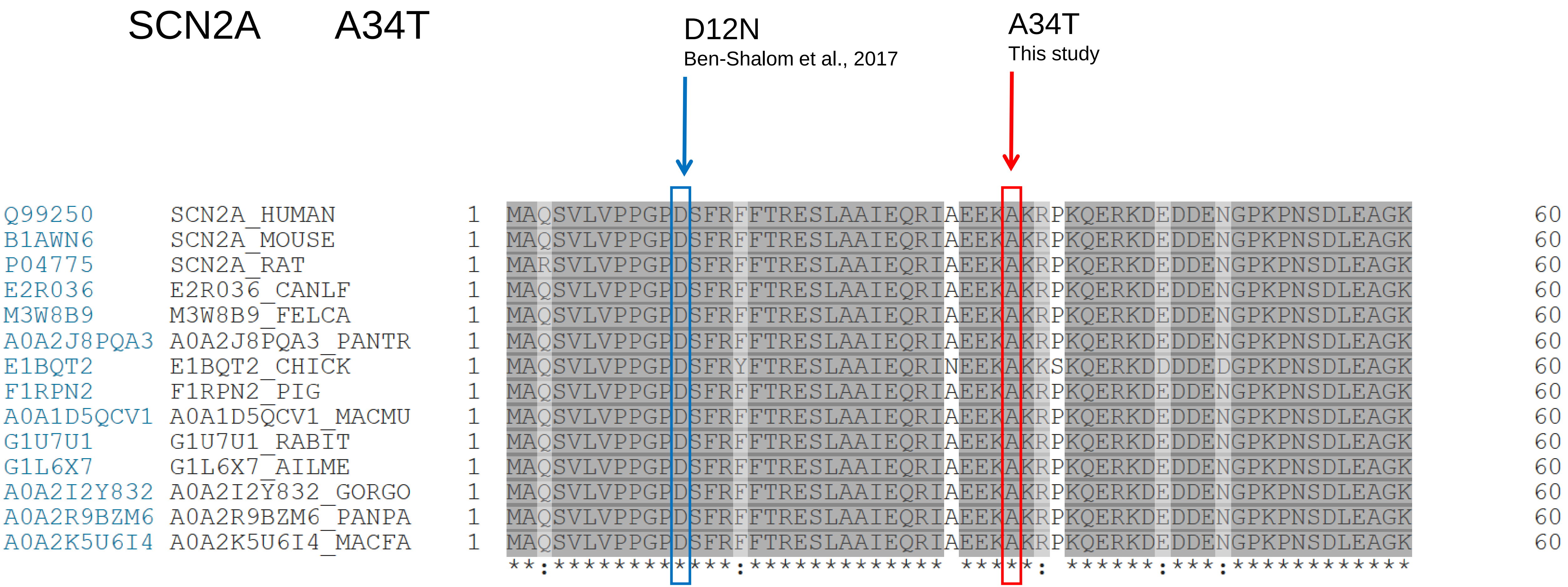


Supplementary Fig.10

**Supplementary Figure 10. Multiple sequence alignment of SCN2A proteins for different  species**

We identified a mutation of SCN2A (A34T) in the family n°23  linked to *FEZF2* allele.

This mutation occurs in a region conserved from chick to primates including humans.

Multiple sequence alignment of SCN2A protein sequences from Human, mouse,rat, dog, cat, chimpanzee, chick, pig, Macaca mulatta, rabbit, Giant panda, gorilla, Bonobo and Macaca fascicularis.

The mutation D12N of the same region (N-terminal cytoplasmic tail) identified by Ben-Shalom et al., 2017 is indicated.

### ERN1 S536L

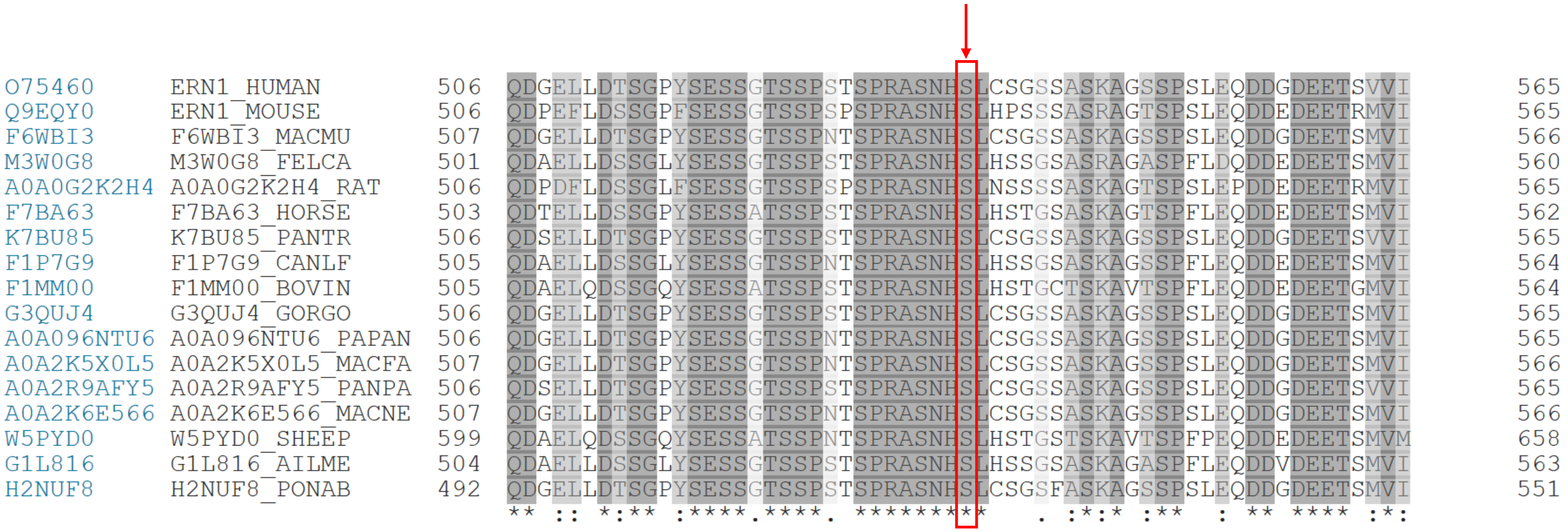


Supplementary Fig.11

**Supplementary Figure 11. Multiple sequence alignment of ERN1 proteins for different species**

We identified a mutation of ERN1 (S536L) in the family n°2 linked to *FEZF2* allele.

This mutation occurs in a region conserved from zebrafish to primates including humans.

Multiple sequence alignment of DIP2A protein sequences from Human, mouse, Macaca mulatta, cat, rat, horse, chimpanzee, dog, Bos taurus, gorilla, Papio anubis, Macaca fascicularis, Bonobo, Macaca nemestrina, sheep, Giant panda and Sumatran orangutan.

### STRIP2 I308V

```
Q9ULQ0      STRP2_HUMAN       291  KVQKRAELGLPPLAEDSIQVVKSMRAASPPSYTLDLGESQLAPPPSKLR-----------  339
Q8C9H6      STRP2_MOUSE       301  KIQKRAELGLPPLAEDSIQVVKSMRAASPPSYTLDLGESQLAPPPSKLR-----------  349
E2RJF2      E2RJF2_CANLF      291  KVQKRAELGLPPLAEDSIQVVKSMRAASPPSYTLDLGESQLAPPPSKLR-----------  339
F6TAH2      F6TAH2_MACMU      291  KVQKRAELGLPPLAEDSIQVVKSMRAASPPSYTLDLGESQLAPPPSKLR-----------  339
M3WSE1      M3WSE1_FELCA      291  KVQKRAELGLPPLAEDSIQVVKSMRAASPPSYTLDLGESQLAPPPSKLR-----------  339
E1BIK9      E1BIK9_BOVIN      291  KVQKRAELGLPPLAEDSIQVVKSMRAASPPSYTLDLGESQLAPPPSKLR-----------  339
E7F9X6      E7F9X6_DANRE      256  KVRMRDHLNLPPLPEDSIKVVRNMRAASPPASAMELIEQQQQQKRGRRSRRSAFVDSLEG  315
G3RKX3      G3RKX3_GORGO      291  KVQKRAELGLPPLAEDSIQVVKSMRAASPPSYTLDLGESQLAPPPSKLR-----------  339
A0A1D5PII0  A0A1D5PII0_CHICK  293  KVRRREELGLPPLPEDSIQVMRSMRAASPPTCSIELAEQQQK------R-----------  335
G1LFF5      G1LFF5_AILME      275  KVQKRAELGLPPLTEDSIQVVKSMRAASPPSYTLDLGESQLAPPPSKLR-----------  323
H2PNH6      H2PNH6_PONAB      291  KVQKRAELGLPPLAEDSIQVVKSMRAASPPSYTLDLGESQLAPPPSKLR-----------  339
A0A096NF25  A0A096NF25_PAPAN  291  KVQKRAELGLPPLAEDSIQVVKSMRAASPPSYTLDLGESQLAPPPSKLR-----------  339
A0A3Q2HTG7  A0A3Q2HTG7_HORSE  289  KVQKRAELGLPPLAEDSIQVVKSMRAASPPSYTLDLGESQLAPPPSKLR-----------  337
A0A2K6BFG4  A0A2K6BFG4_MACNE  291  KVQKRAELGLPPLAEDSIQVVKSMRAASPPSYTLDLGESQLAPPPSKLR-----------  339
H2QVC9      H2QVC9_PANTR      291  KVQKRAELGLPPLAEDSIQVVKSMRAASPPSYTLDLGESQLAPPPSKLR-----------  339
A0A2R9B940  A0A2R9B940_PANPA  288  KVQKRAELGLPPLAEDSIQVVKSMRAASPPSYTLDLGESQLAPPPSKLR-----------  336
                                   *:: * .*.**** ****:*::.*******: :::* *.*
```

Supplementary Fig.12

**Supplementary Figure 12. Multiple sequence alignment of STRIP2 proteins for different species**

We identified a mutation of STRIP2 (I308V) in the family n°6 linked to *FEZF2* allele.

This mutation occurs in a region conserved from zebrafish to primates including humans.

Multiple sequence alignment of DIP2A protein sequences from Human, mouse, dog, Macaca mulatta, cat, Bus Taurus, zebrafish, gorilla, chick, Giant panda, Sumatran orangutan, Papio anubis, horse, Macaca nemestrina, chimpanzee and Bonobo.

Supplementary Figure 13

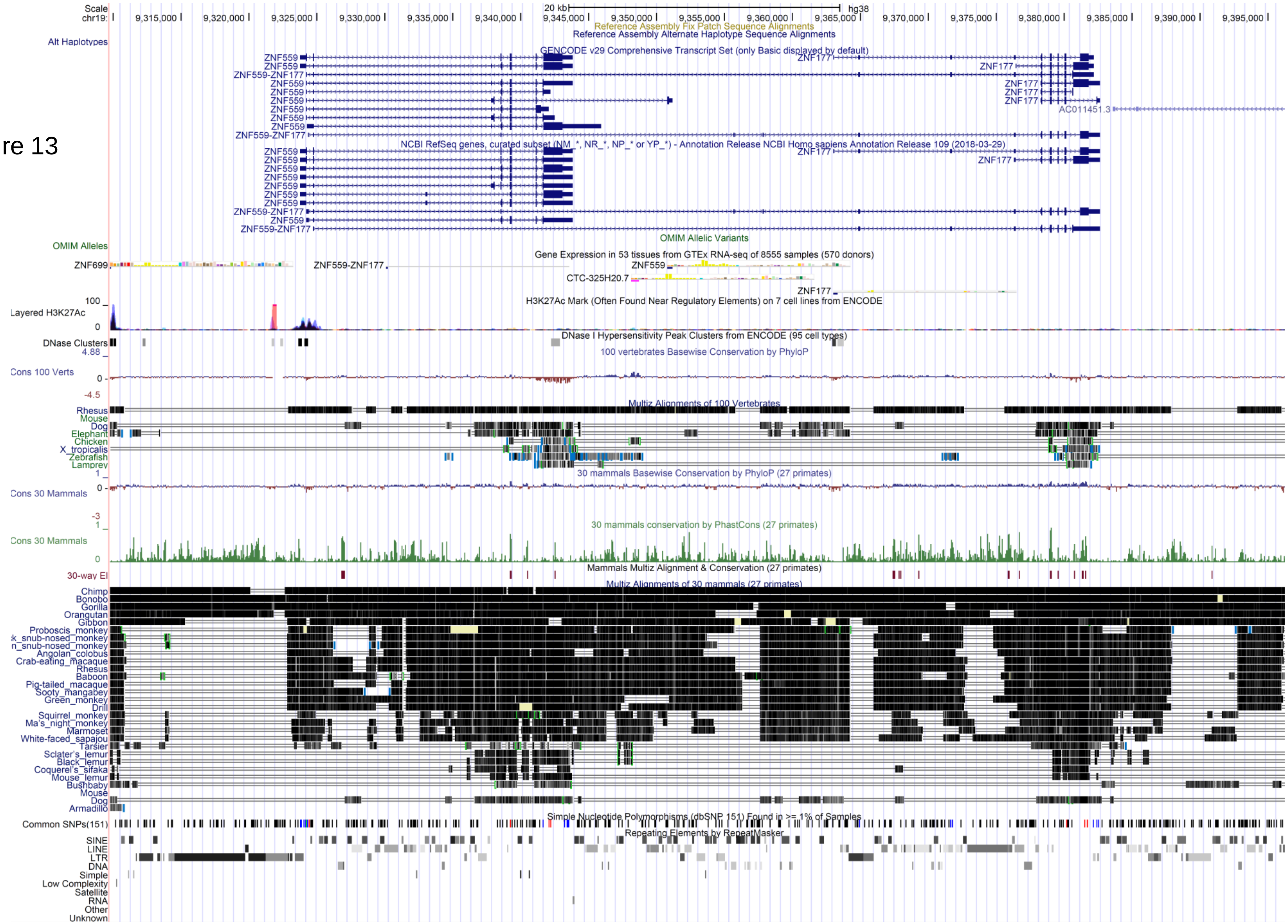

**Supplementary Figure 13. *ZNF559* gene in human genome with its primate-specific conservation**

Analysis of *ZNF559* in human genome (human HG38). Note that it is fully conserved in primates but not in other mammals, suggesting that *ZNF559* is a primate-specific innovation.

### Supplementary Figure 14

A

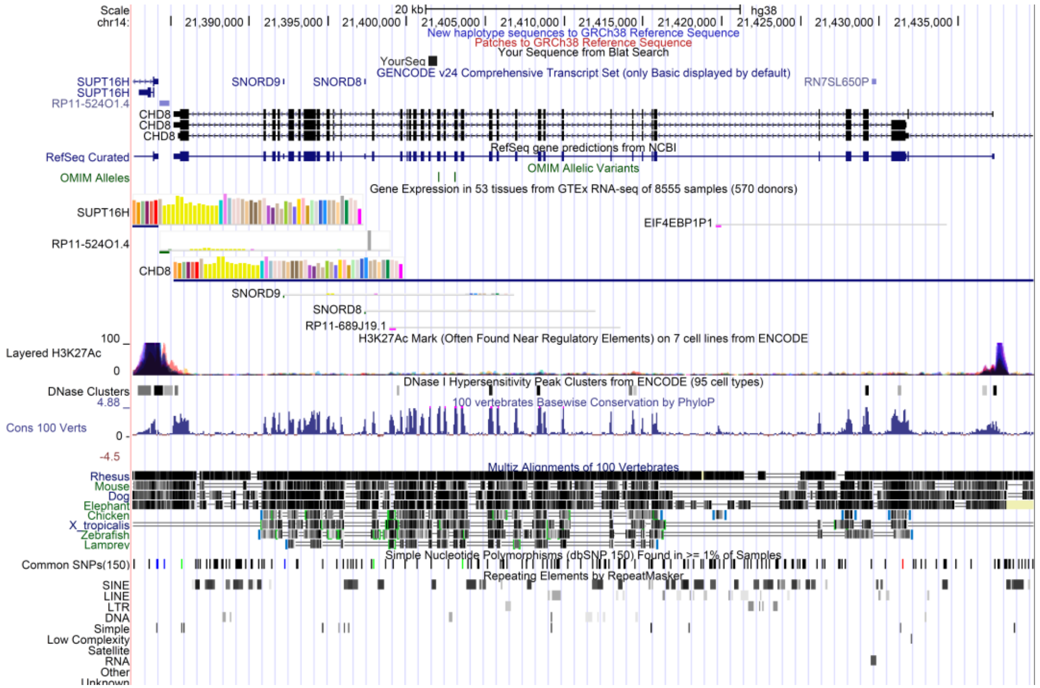


B

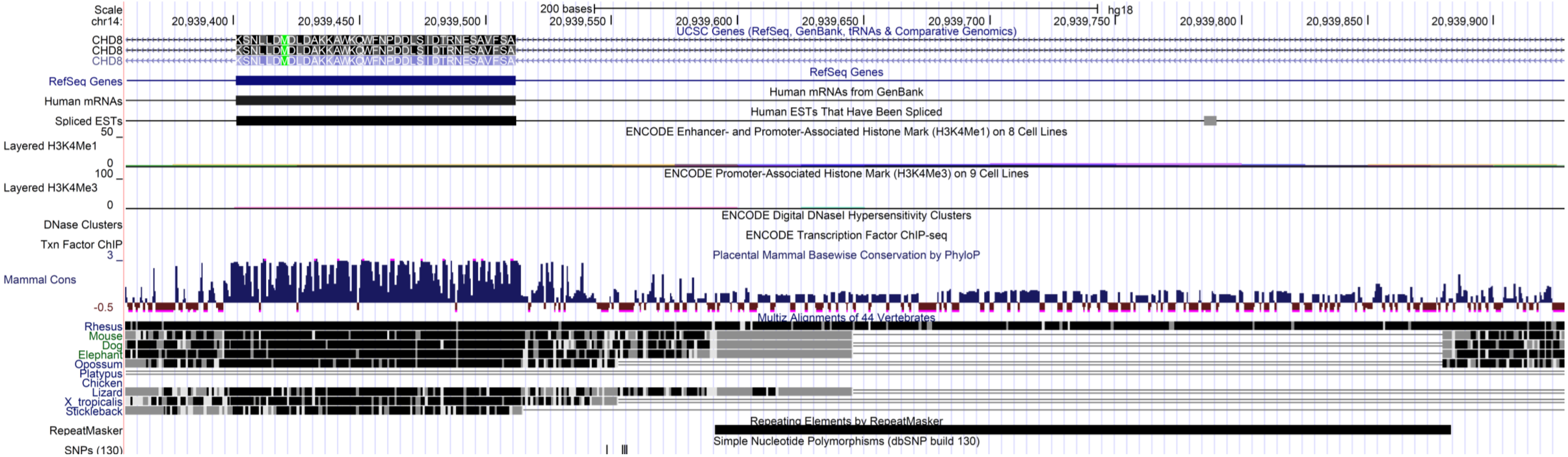

**Supplementary Figure 14. Localization of the deletion in *CHD8* gene identified in a FEZF2 allele-unlinked family.**

We identified a deletion in the *CHD8* gene in *FEZF2* allele-unlinked family 20. This mutation involves the full deletion of a *CHD8* exon that follows the helicase domain. (**A**). Localization of the 571bp deletion of the *CHD8* gene locus. (**B**) Deletion of 571bp indicating the *CDH8* exon involved and the phylogenetically conserved region.

Supplementary Figure 15

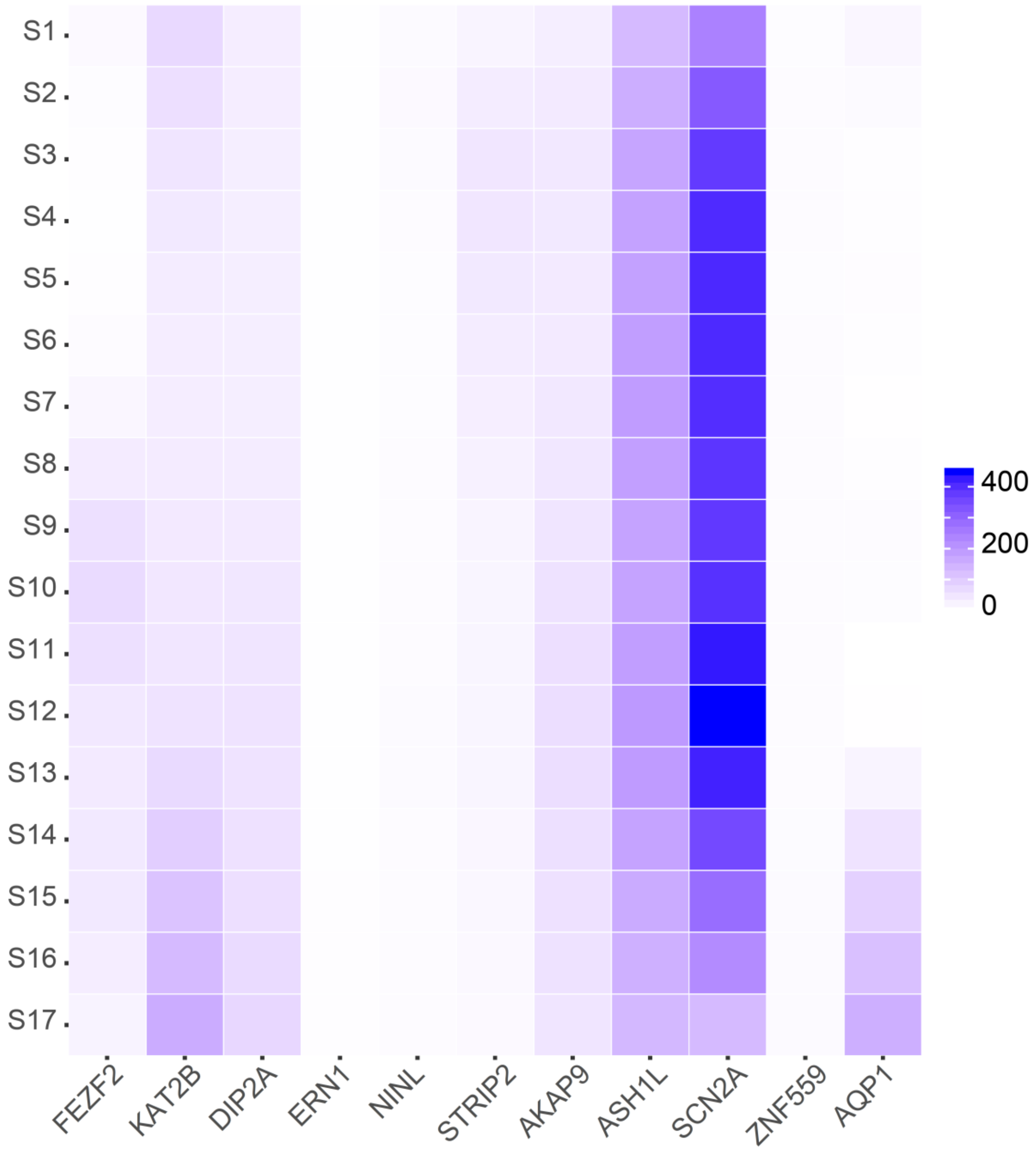

**Supplementary Figure 15. Heatmap showing gene expression in RPKM across 17 prefrontal cortical sections.**

We used data from adult human brain (He et al. 2017) for *FEZF2* and nine other ASD associated genes.
